# Unraveling the role of *Xist* in X chromosome inactivation: insights from rabbit model and deletion analysis of exons and repeat A

**DOI:** 10.1101/2023.09.21.558801

**Authors:** Mingming Liang, Lichao Zhang, Liangxue Lai, Zhanjun Li

**Affiliations:** Key Laboratory of Zoonosis Research, Ministry of Education, Jilin University, Changchun 130062, China; China-New Zealand Joint Laboratory on Biomedicine and Health, CAS Key Laboratory of Regenerative Biology, Guangdong Provincial Key Laboratory of Stem Cell and Regenerative Medicine, Centre for Regenerative Medicine and Health, Hong Kong Institute of Science and Innovation, Guangzhou Institutes of Biomedicine and Health, Chinese Academy of Sciences, Guangzhou 510530, China; Research Unit of Generation of Large Animal Disease Models, Chinese Academy of Medical Sciences (2019RU015), Guangzhou, 510530, China

**Author notes:** Corresponding author.; (Zhanjun Li); (Liangxue Lai).

## Abstract

X chromosome inactivation (XCI) is a dosage compensation process that aims to equalize the expression of X-linked genes between males and females. Initiation of XCI relies on *Xist*, which is continuously expressed in somatic cells during XCI maintenance. How *Xist* impacts XCI maintenance remains unclear, and its functional motifs remain unclear. In this study, we conducted a comprehensive analysis of the *Xist*. The results demonstrate that rabbits serve as an ideal non-primate animal model for investigating the functions of *Xist* in vivo. And homozygous knockout of exon 1, exon 6 and repeat A in females resulted in embryonic lethality. However, X^ΔReA^X females with *Xist* expression from the intact X chromosome, did not display any abnormalities. Interestingly, there were no significant differences between females with homozygous knockout of exons 2-5 and wild-type rabbits. This suggests that exons 2, 3, 4, and 5 of *Xist* are less important for XCI. These insights provide valuable knowledge about the functional mechanism of *Xist*.

## Introduction

X chromosome inactivation (XCI) is a dosage compensation mechanism that has evolved in marsupial and placental mammals to equalize the level of X-linked gene expression between females (XX) and males (XY) (Lyon, 1961, 1962; Graves, 1996; Arthold *et al*, 2011; Morey & Avner, 2011). The initiation of XCI is genetically controlled by a master regulatory locus named the X-inactivation center (Xic) (Avner & Heard, 2001; Brockdorff & Duthie, 1998a; Goto & Monk, 1998a; Heard *et al*, 1997a; Willard, 1996). In mice and humans, dosage compensation is mediated by a long noncoding RNA (lncRNA) termed *Xist* (Brown *et al*, 1992; Brockdorff *et al*, 1992), which is up-regulated from one of the two X chromosomes. Its RNA accumulates over the inactive X chromosome (Xi) in cis to trigger gene silencing (Brockdorff & Duthie, 1998b; Heard *et al*, 1997b; Lyon, 1999). Once established, XCI is stably inherited upon successive cell divisions in female somatic cells. Extensive studies have confirmed that *Xist* is both necessary and sufficient for XCI (Loda *et al*, 2022; Pandya-Jones *et al*, 2020a; Chu *et al*, 2015).

Since the proposal of XCI by Lyon in 1961 (Lyon, 1961), mouse models have been widely used to investigate the molecular mechanisms underlying X inactivation (Morey & Avner, 2011; Avner & Heard, 2001; Brockdorff & Duthie, 1998b; Dossin *et al*, 2020; Arnold, 2022; Markaki *et al*, 2021; Yang *et al*, 2011). In mice, the *Xist* gene is composed of seven exons, which are interspersed with repetitive sequences known as repeats A to F (Brockdorff *et al*, 1992). These exons and repetitive sequences play crucial roles in the localization and spreading of *Xist* RNA along the inactive X chromosome (Xi) (Loda & Heard, 2019). Repeat A, in particular, has been extensively studied and found to be essential for *Xist*-mediated gene silencing. It also facilitates the recruitment of chromatin-modifying factors through interactions with specific proteins (Brockdorff, 2018; Bousard *et al*, 2019a; Carter *et al*, 2020; Colognori *et al*, 2020a; Trotman *et al*, 2020; Lu *et al*, 2016; Royce-Tolland *et al*, 2010; Wutz *et al*, 2002; Jones *et al*, 2022; Sarkar *et al*, 2015). Other repeats, such as B and C, also contribute to *Xist*-mediated gene silencing and the establishment of the inactive chromatin state (Pintacuda *et al*, 2017; Nakamoto *et al*, 2020; Bousard *et al*, 2019b; da Rocha *et al*, 2014; Colognori *et al*, 2020b, 2019; Wei *et al*, 2021). Repeat E is crucial for the proper localization of *Xist* RNA on the inactive X chromosome (Xi) and facilitates gene silencing (Pandya-Jones *et al*, 2020c; Ridings-Figueroa *et al*, 2017; Sunwoo *et al*, 2017; Smola *et al*, 2016; Yue *et al*, 2017; Cherney *et al*, 2023; Pandya-Jones *et al*, 2020b). Studies conducted on mouse embryonic stem cells have identified the importance of specific exons in XCI. For instance, the 5’ region of *Xist* plays a critical role in gene silencing (Trotman *et al*, 2020; Coker *et al*, 2020), while exon 4 exhibits a lesser impact on X inactivation (Caparros *et al*, 2002). In human cells, different exons, such as exon 5, are vital for maintaining XCI, while exons 2, 3, and 4 are relatively less significant (Lee *et al*, 2019). These findings have been demonstrated at the cellular level, which may not fully to mimic the XCI in vivo.

A significant amount of our current understanding of the mechanisms of XCI comes from studies conducted on mice and in vitro female stem cells (Berletch *et al*, 2015; Morey & Avner, 2011; Yamada *et al*, 2015). *Xist*, a gene involved in XCI, shows species-specific differences in its regulation and function. In humans, *XIST* is expressed on both X chromosomes, which undergo random XCI during cell differentiation (Briggs *et al*, 2015; Okamoto *et al*, 2011). On the other hand, mice exhibit paternal *Xist* expression during the initial stage of X chromosome inactivation, followed by random X chromosome inactivation during the subsequent stage. Therefore, there are notable differences between the mechanisms of X inactivation in mice and humans(Okamoto *et al*, 2011). Consequently, it is crucial to establish suitable animal models for investigating the functions of *XIST* and studying the mechanisms of XCI.

According to a study by Okamoto et al. published in Nature, rabbits exhibit a similar XCI mechanism to humans during early embryogenesis(Okamoto *et al*, 2011). Both species undergo random X chromosome inactivation in both stages of *XIST* expression. In contrast, the house mouse displays paternal *Xist* expression during the initial stage of X chromosome inactivation, followed by random X chromosome inactivation during the subsequent stage (Fig 1D). These findings highlight the differences in the XCI mechanism in regulating *Xist* expression in mice and humans, and the rabbit would be an ideal animal model for studying XCI.

**Figure 1.**
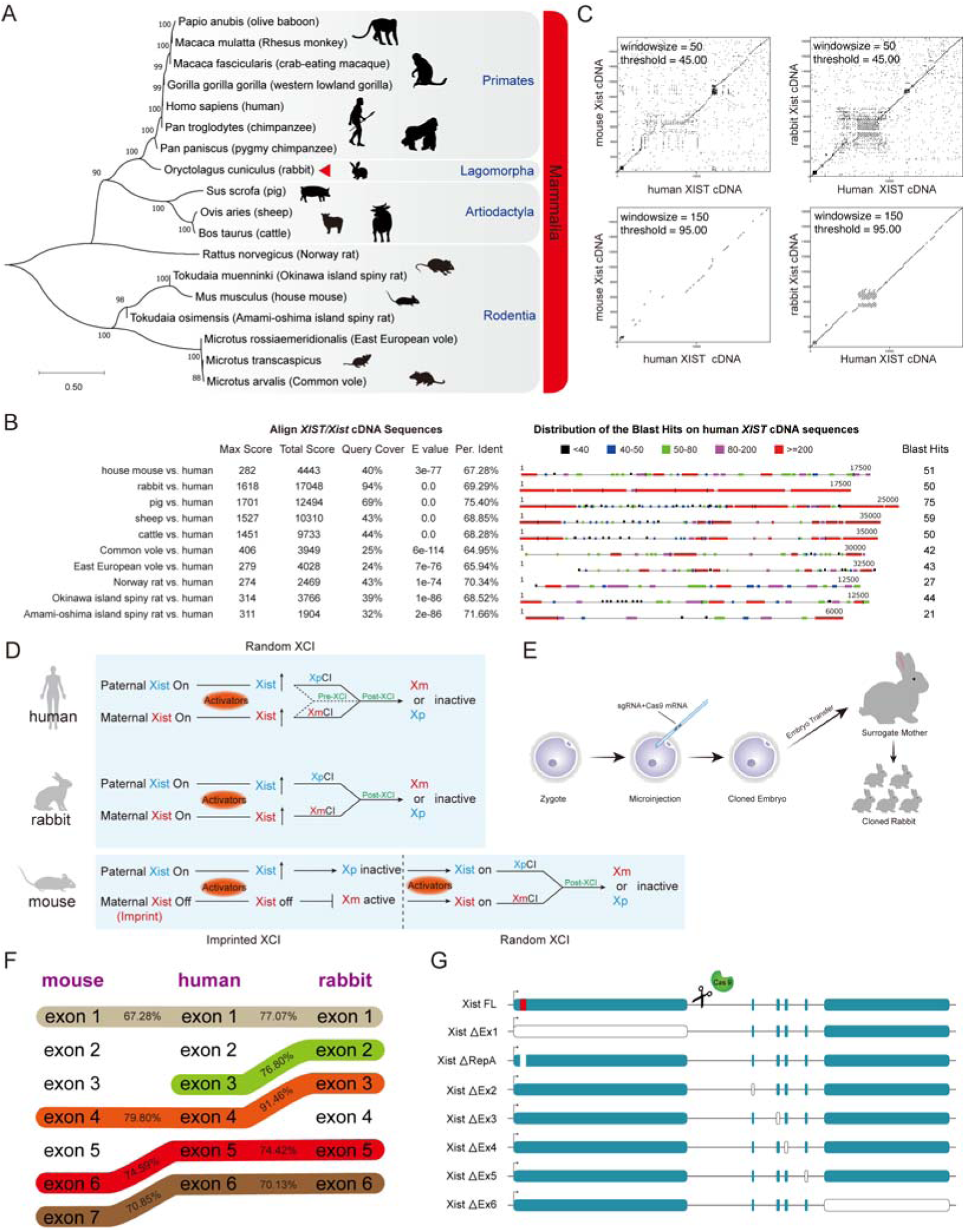
Comparison of *Xist* between different species. (**A**) Evolutionary relationships of the *XIST/Xist* gene among Primates, Lagomorpha, Artiodactyla, and Rodentia. Mammalian phylogeny was estimated using maximum likelihood from 18 nucleotide sequences. Clades discussed in the text are labeled. Bootstrap support values ≥88% are indicated at nodes. The scale bar indicates evolutionary distance. (**B**) *Xist* sequences were aligned to humans using the NCBI BLAST server. (**C**) Dot plot analysis of *Xist/XIST* cDNA sequences in mouse, rabbit, and human using the EMBOSS dot-matcher program. (**D**) Hypothesis explaining differences in *Xist/XIST* regulation and XCI initiation observed in mouse, rabbit, and human embryos, based on previous work (Okamoto *et al*, 2011). (**E**) Schematic diagram illustrating the establishment of a cloned rabbit model using standard microinjection procedures. (**F**) Schematic representation of the sequence homology of *Xist/XIST* exons across mice, humans, and rabbits. The percent identity mentioned in the text is indicated. (**G**) Schematic representation of the *Xist* knock-out created in rabbits using CRISPR/Cas9; the different exons are highlighted in colored boxes, with repeat A highlighted in the red box. Deleted exons are represented as white boxes.

**Figure EV1.**
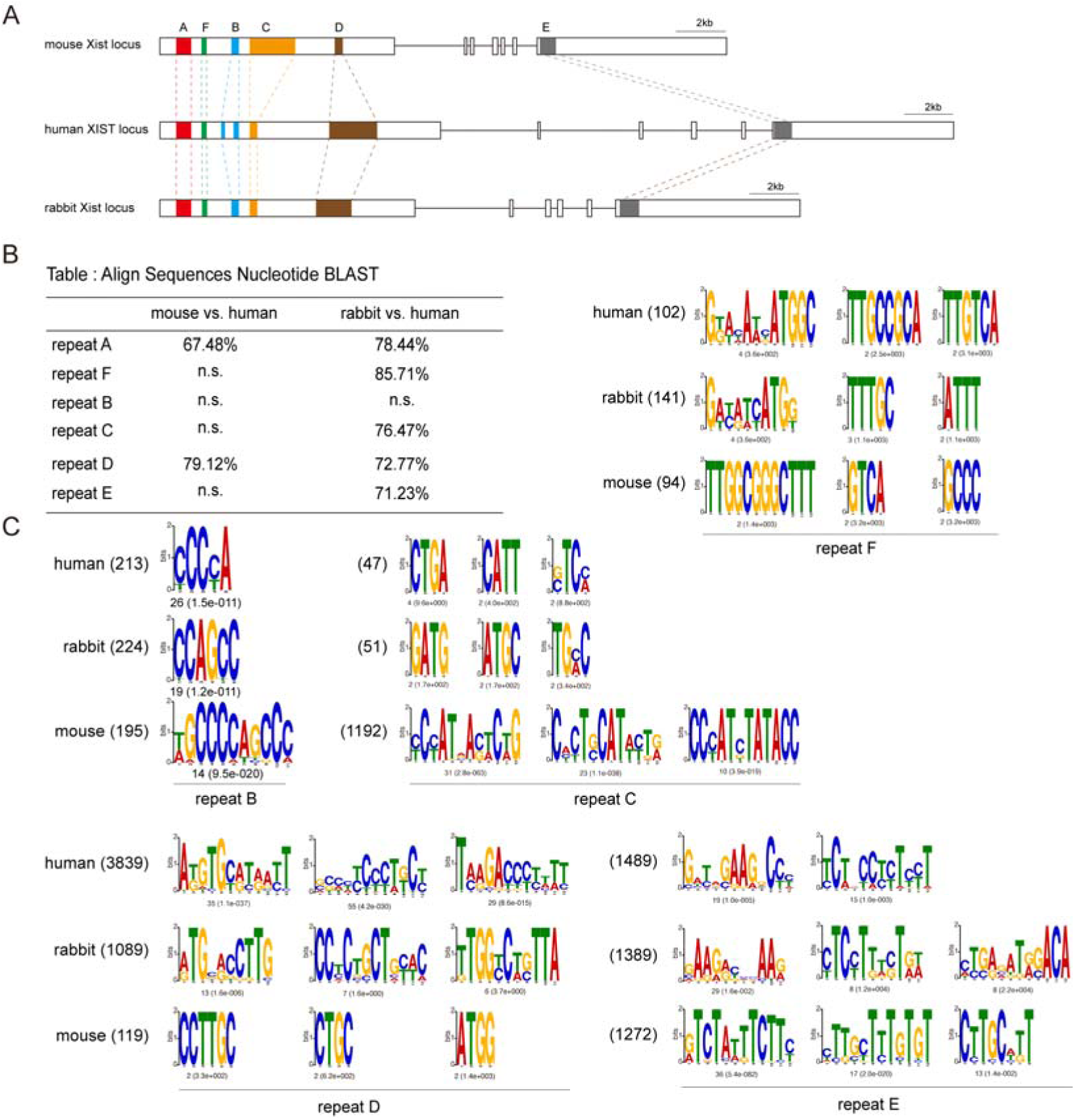
Characterization of *Xist*/*XIST* repeat sequences. (**A**) Comparison of the gene structure and tandem repeats of *Xist*/*XIST* in mouse, human, and rabbit. The core repeat regions are highlighted in different colors. (**B**) The BLAST analysis shows the nucleotide sequence homology of *Xist*/*XIST* repeat among mouse, human, and rabbit, as indicated in the table. (**C**) Comparison of the top three motifs identified by MEME in *Xist*/*XIST* repeat A. The number of sites and E-value for each motif logo are provided. The length of each repeat is mentioned in nucleotides (shown in parentheses). All *XIST/Xist* sequences used in this study are listed in Appendix Table 4-6.

In this study, we conducted a comprehensive phylogenetic analysis which revealed a close relationship between rabbits and primates, and also showed that human *XIST* exhibited a higher sequence homology to rabbits than mice, suggest that rabbits would be an ideal animal model for studying the XCI mechanism. In addition, the functional of *Xist* RNA transcript was determined by deleting exon 1-6 and repeat A of rabbit *Xist* using CRISPR/Cas9. Overall, these findings contribute to a deeper understanding of the functional mechanisms underlying *Xist*-induced XCI in animal level.

## Results

### Rabbits are the ideal non-primate animal model for studying *Xist* in vivo

To investigate the functional role of *Xist*, we conducted a phylogenetic analysis using Molecular Evolutionary Genetics Analysis software (MEGA11) and employed the maximum composite likelihood method with 1000 bootstrap replicates (Tamura *et al*, 2021). Reference sequences of the *Xist* gene were obtained from the NCBI database (Sayers *et al*, 2021). The phylogenetic tree analysis of the *XIST/Xist* gene demonstrated a close relationship between rabbit species and humans, while excluding non-human primates (Fig 1A). The three species, namely pigs, cows, and sheep, belong to the same sister branch in the taxonomic status of the evolutionary tree, indicating a close evolutionary relationship. The results revealed that rabbits and primates are part of the same main branch of the phylogenetic tree, indicating a closer relationship of the *XIST/Xist* gene between rabbits and primates. This finding positions rabbits as an ideal non-primate animal model for studying *Xist* in vivo.

Additionally, we conducted a comparative analysis of *Xist* DNA sequence homology across various non-primate species. The results demonstrated that rabbits exhibited the highest overall score and sequence coverage, scoring a total of 17048 and achieving a coverage score of 94%. In comparison, the house mouse, which is a classic animal model, showed a total score of 4443 and a coverage of 40%, indicating a low overall DNA homology to human *Xist*, suggesting the higher homology of rabbit *Xist* to humans (Fig 1B). To further assess the similarity of *Xist* sequences between house mouse, rabbit and human, we conducted a Dot-Plot analysis (Madeira *et al*, 2022). Consistent results were observed across different window sizes and thresholds, confirming that rabbits have higher sequence similarity to humans than to mice (Fig 1C).

In addition, rabbits exhibit a similar XCI mechanism to humans during early embryogenesis(Okamoto *et al*, 2011). Humans and rabbits undergo random X chromosome inactivation in both stages of *XIST* expression. In contrast, mice display paternal *Xist* expression during the initial stage of X chromosome inactivation, followed by random X chromosome inactivation during the subsequent stage (Fig 1D).

Then, we conducted a comparative analysis to assess the homology of exons among mice, rabbits, and humans *XIST/Xist* (Figs 1F and EV2A). The results showed that humans *XIST* exon 1 had a higher homology of 77.07% with rabbits *Xist* exon 1, compared to the 67.28% homology observed with mice *Xist* exon 1. Interestingly, we did not find any homologous sequence of humans *XIST* exon 2 in both mice and rabbits. On the other hand, humans *XIST* exon 3 showed a homology of 76.80% with rabbits *Xist* exon 2, but no homologous sequence was detected in the mice. Furthermore, humans *XIST* exon 4 had a higher homology of 91.46% with rabbits *Xist* exon 3, compared to 79.80% homology with mice *Xist* exon 4. Additionally, humans *XIST* exon 5 demonstrated homologies of 74.59% and 74.42% with mice *Xist* exon 6 and rabbits *Xist* exon 5, respectively. Similarly, humans *XIST* exon 6 showed homologies of 70.85% and 70.13% with mice *Xist* exon 7 and rabbits *Xist* exon 6, respectively.

Besides, we also conducted a comparison of the similarity of other repeat sequences on the *Xist* loci of human, mouse, and rabbit (Fig EV1A). We first evaluated the homology of the different repeat sequences and found that the homology between rabbit repeat F and human repeat F was 85.71%. However, there was no corresponding homologous sequence of human repeat F in mice. Similarly, the homology between rabbit repeat C and human repeat C was 76.47%, but no corresponding homologous sequence was found in mice. The homology between rabbit repeat D and human repeat D was 72.77%, while mouse repeat D exhibited 79.12% homology to its human counterpart. Rabbit repeat E exhibited a 71.23% homology to the human sequence, while no corresponding homologous sequence was found in the mouse (Fig EV1B). Furthermore, we compared the motif similarity of the different repeat sequences and observed that the rabbit repeats were more similar to human repeats in terms of length (Fig EV1C). Notably, repeat B contained repeat units “enriched in cytosine bases”, and repeat E showed greater similarity between rabbit and human sequences, whereas the mouse sequence exclusively consisted of repeat units “enriched in thymine bases”.

In summary, these findings solidify the importance of rabbits as an invaluable model for understanding the functions of *Xist*. To generate cloned animals, we co-injected Cas9 mRNA and sgRNA into one-cell stage embryos and transferred them into surrogate mother rabbits (Figs 1E and EV2B). We utilized CRISPR/Cas9 technology to disrupt the exon sequences of *Xist* in rabbits (Sui *et al*, 2016; Yao *et al*, 2020), specifically targeting exons 1-6 and repeat A, to investigate the functional role of the *Xist* (Fig 1G).

### Deletion of exon 1 in female rabbits does not survive

Sequence homology analysis was performed on *Xist* exon 1 in mice, humans, and rabbits using the NCBI BLAST service. The coverage rate of rabbit *Xist* exon 1 was determined to be 95% when compared to the human *XIST* sequence, which was significantly higher than the 41% observed in mice. Moreover, the percentage identity between rabbit *Xist* exon 1 and human *XIST* exon 1 was found to be 77.07%, while the percentage identity between mice and humans was 67.28% (Fig 2A). Dot-Plot analysis confirmed that rabbit *Xist* exon 1 exhibited higher homology to human compared to mouse (Fig 2B).

**Figure 2.**
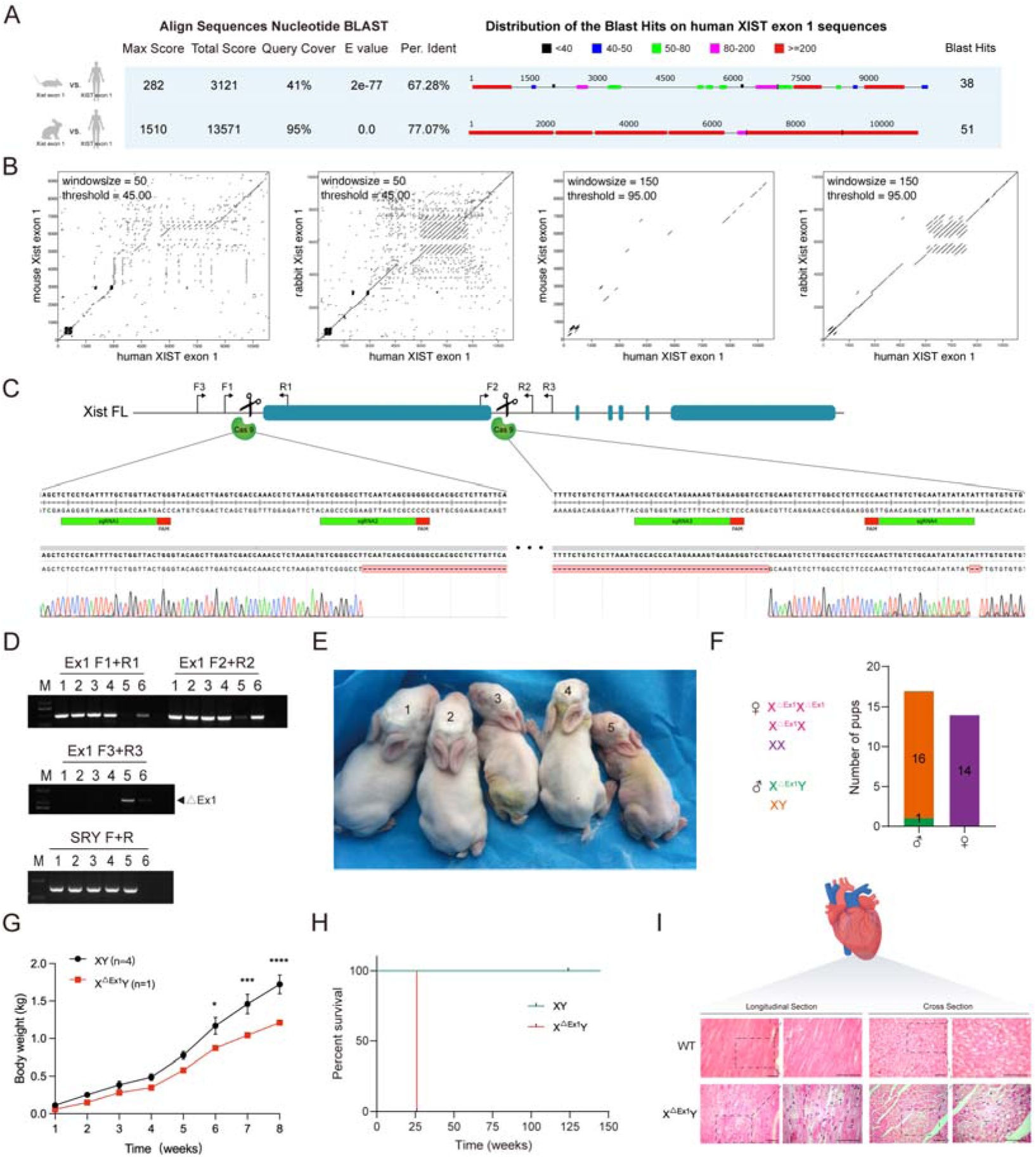
Male rabbit with *Xist* exon 1 deletion exhibit a loose arrangement of the myocardial fibers. (**A**) The BLAST results of *Xist/XIST* exon 1 are compared between mouse, human, and rabbit. (**B**) Dot plot analysis of the *Xist/XIST* exon 1 sequence in mouse, rabbit, and human. (**C**) Target loci and Sanger sequencing results demonstrate the knock-out of *Xist* exon 1 in F0 male rabbits. All sgRNA sequences are listed in Appendix Table 1. (**D**) Founder rabbits from the F0 generation are identified through agarose gel electrophoresis. (**E**) The gross appearance of rabbits from the F0 generation at day 7 reveals that *Xist* exon 1 knock-out rabbits exhibit developmental delay. (**F, left**) Schematic for generating heterozygous *Xist* exon 1 deletants. (**Right**) Genotype data for F0; the number of pups for each genotype is listed. (**G**) The body weight of male X^△Ex1^Y F0 rabbit and male littermate controls (n = 4). Error bars indicate mean ± SEM. (**H**) The survival curve for X^△Ex1^Y and male littermate controls. (**I**) H&E staining for cardiac muscle from X^△Ex1^Y and male control animal. Scale bars, 50um.

**Figure EV2.**
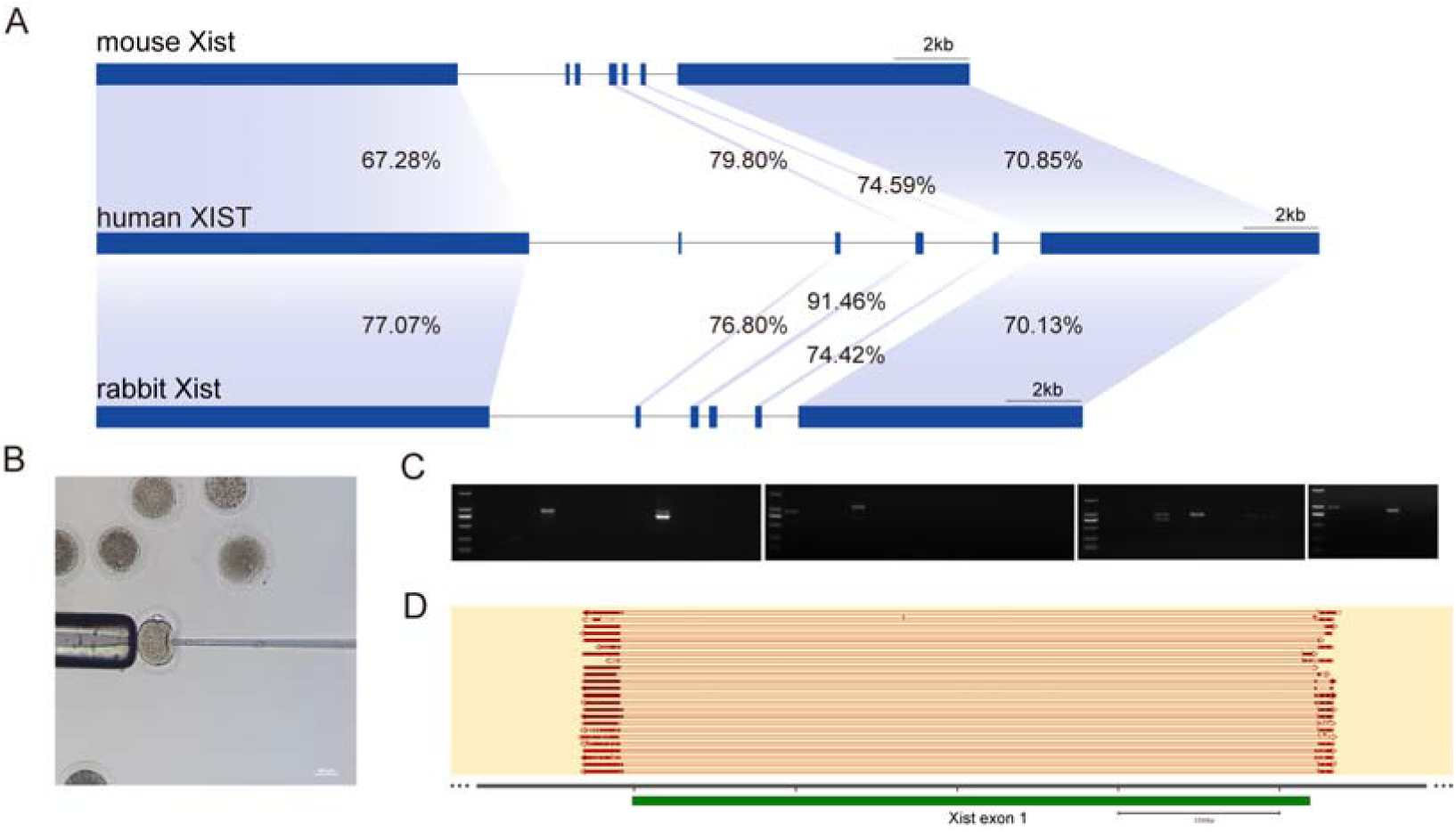
Characterization of the *Xist* exon 1 knock-out mutant in the embryo. (**A**) Comparison of the gene structure and homology region of exons between mouse, rabbit, and human *Xist*/*XIST*. Exons are marked in dark blue. (**B**) Diagram illustrating microinjection in a rabbit embryo. (**C**) Identification of the cloned embryo through agarose gel electrophoresis using primers F3/R3. (**D**) Characterization of the *Xist* exon 1 knock-out mutant in the cloned embryo. Scale bars: 2500bp. All sgRNA sequences are listed in Appendix Table 1.

To generate knock-out rabbits using CRISPR/Cas9 technology, we designed four single-guide RNAs (sgRNAs) targeting exon 1 of *Xist*. Genotyping was performed through PCR using four sets of primers specifically designed to target certain genes, including the Sry gene, which is exclusively present in males, and the Sanger sequencing was conducted to confirm the genotyping results (Fig 2C and D). No homozygous knockout female rabbits were identified among the genotyping results of the founder animals (Fig 2F). However, we were able to successfully obtain a hemizygous knockout male rabbit, referred to as #5 (Fig 2E). Regrettably, this male rabbit showed developmental delays (Fig 2G) and eventually succumbed at 25 weeks (Fig 2H). Upon examination of the deceased individual’s myocardium, a loose arrangement of cardiac fibers (Fig 2I) was observed in the #5 rabbit.

Given the consistent challenges encountered in obtaining homozygous knockout female individuals, we hypothesized that the X^ΔEx1^X^ΔEx1^ homozygous mutant females could be generated but would die early in embryogenesis. To investigate this possibility, we conducted an embryonic-level investigation by employing a fibroblast injection method to introduce sgRNA and Cas9 RNA into zygotes. After a week-long incubation period, genotyping was performed using PCR and Sanger sequencing. Remarkably, the results revealed successful large-scale deletions at the embryonic level (Fig EV2C and D). The results showed that the X^ΔEx1^X^ΔEx1^ homozygous mutant females can be generated but perish early in embryogenesis.

### Deletion of *Xist* repeat A in rabbits results in embryonic lethality

To investigate the impact of *Xist* repeat A on individual development, we conducted a comparative analysis of sequence homology in mice, humans, and rabbits. Interestingly, our findings showed a higher level of homology (78.44%) between rabbit and human repeat A sequences, while the homology between mouse and human sequences was 67.48% (Fig 3A), and the sequence of rabbit repeat A bore a closer resemblance to the human (Fig EV3A), which was also confirmed by using Dot-Plot analysis (Fig 3B). In addition, the more similar of repeat motifs were determined in rabbit and human (Figs 3C and EV3B). Furthermore, the consistent major stem-loop structures in both rabbit and human repeat A, but the mouse sequence showed discrepancies (Fig 3D). Specifically, the RNA hairpin in rabbit and human sequences consisted of 12 nucleotides is AUCG tetraloop, whereas AWCG tetraloop in mice. These results clearly demonstrate that the rabbit *Xist* repeat A sequence is more similar to the human sequence. Subsequently, two pairs of single guide RNAs (sgRNAs) were designed to target and delete the *Xist* repeat A. The genotyping was determined by performing PCR, and the results were further confirmed through Sanger sequencing (Fig 3E). Then the homozygote knockout *Xist* repeat A (X^ΔReA^X^ΔReA^) females were generated by backcross breeding (Figs 3F and EV3C). However, despite multiple attempts, we were unable to generate X^ΔReA^X^ΔReA^ female rabbits (Figs 3F and EV3D). Anatomical data showed that at rabbit embryonic day 12 (E12) we only obtained X^ΔReA^X, X^ΔReA^Y, XY embryos and no X^ΔReA^X^ΔReA^ embryos, indicating that X^ΔReA^X^ΔReA^ could not survive to E12. (Fig 3G).

**Figure 3.**
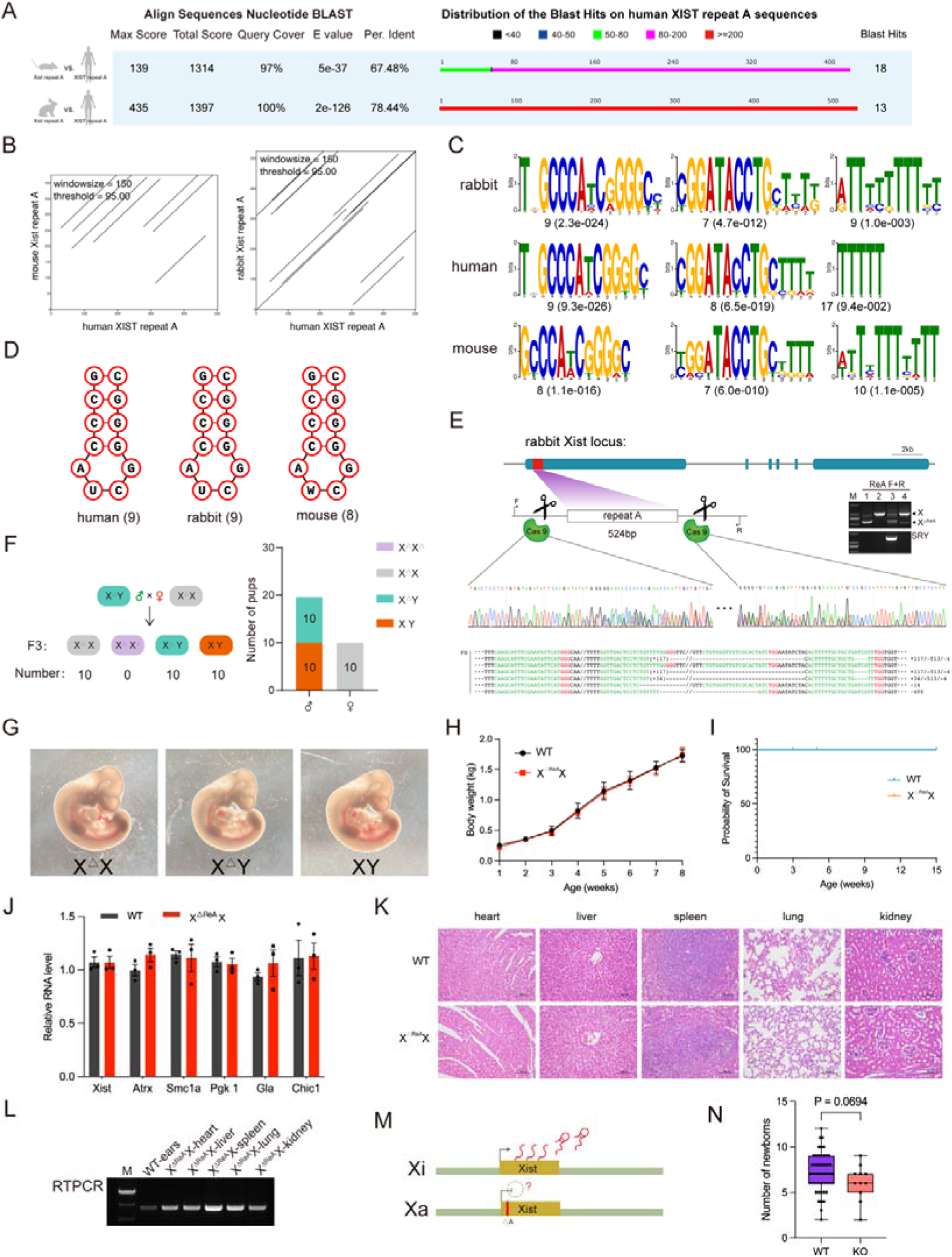
*Xist* repeat A homozygous knockout rabbit does not survive. (**A**) The BLAST result compares the *Xist*/*XIST* repeat A sequences in mouse, human, and rabbit. (**B**) Dot plot analysis illustrates the sequence similarities between mouse, rabbit, and human *Xist*/*XIST* repeat A. (**C**) The top motifs identified by MEME in *Xist*/*XIST* repeat A are compared, with the number of sites and E-value displayed below each motif logo. (**D**) Secondary structure analysis of *Xist*/*XIST* repeat A is presented for human, rabbit, and mouse. The total numbers of stem loops in repeat A are listed. (**E**) Target loci and Sanger sequencing results provide evidence of *Xist* repeat A knock-out in F0 rabbit. All sgRNA sequences are listed in Appendix Table 1. (**F, left**) A schematic depicts the process of generating homozygous *Xist* repeat A deletants. (**Right**) Genotype data for F3; the number of pups for each genotype is listed. (**G**) Phenotypic comparison of WT and mutant littermates at E12. (**H**) Body weight of X^△ReA^X and WT rabbits. Error bars indicate mean ± SEM. (**I**) Survival curve for X^△ReA^X and WT rabbits. (**J**) Expression of X-linked genes in X^△ReA^X and WT rabbits. Error bars indicate mean ± SEM. (**K**) H&E staining for main organs from X^△ReA^X and control animal. Scale bars, 100um. (**L**) Gene expression analysis by RT-PCR. (**M**) A model for *Xist*-mediated transcriptional silencing across the X chromosome in X^△ReA^X rabbits. (**N**) Newborn data from cross-breeding between X^△ReA^X / X^△ReA^Y compared to WT control (box and whiskers plot, min. to max., all points shown).

**Figure EV3.**
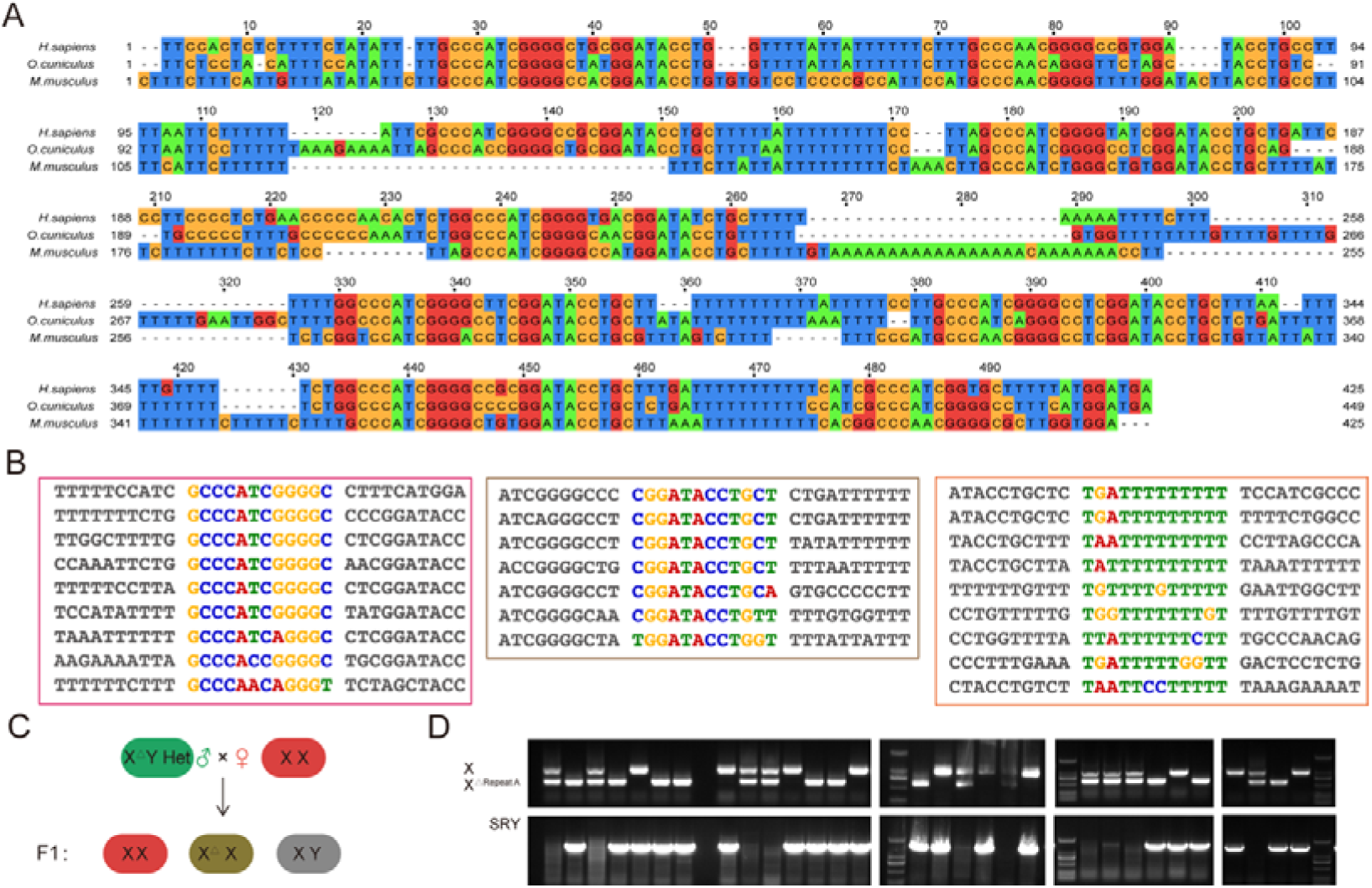
Alignment of tandem repeats in *Xist* repeat A. (**A**) Multiple sequence alignment of repeat A in human, rabbit, and mouse. (**B**) Alignment of three sets of tandem repeats in rabbit *Xist* repeat A. (**C**) Schematic illustrating the generation of the F1 generation. (**D**) Agarose gel electrophoresis result of PCR product in the F3 generation.

However, there are no significant differences were observed in the body weight (Fig 3H) and survival rates (Fig 3I) and X-linked gene expression levels (Fig 3J) of X^ΔReA^X females compared to the WT rabbits. In addition, the normal development of the heart, liver, spleen, lungs and kidneys in X^ΔReA^X females was comparable to that of WT rabbits (Fig 3K).

The RT-PCR results indicated that *Xist* expression was observed in X^ΔReA^X rabbits, which is consistent with *Xist* expression in WT rabbits (Fig 3L). These findings suggest that the *Xist* repeat A is transcribed from the complete X chromosome (Fig 3M). Additionally, the average number of offspring in the X^ΔReA^X female and X^ΔReA^Y male cross-group (6.091 offspring) was lower than that in the WT group (7.118 offspring), further supporting the results that embryos lacking *Xist* repeat A function do not survive (Fig 3N).

### Deletions of *Xist* exon 2 in rabbits are viable and develop normally

It is shown that there no homologous sequence of human *XIST* exon 3 in mice. However, there is a higher homology (76.80%) between rabbit *Xist* exon 2 and human *XIST* exon 3 (Fig 4A), which was also confirmed by Dot plot analysis (Fig EV4A). Thus, the deletions of *Xist* exon 2 in rabbits were used to mimic the function of *XIST* exon 3 in humans.

**Figure 4.**
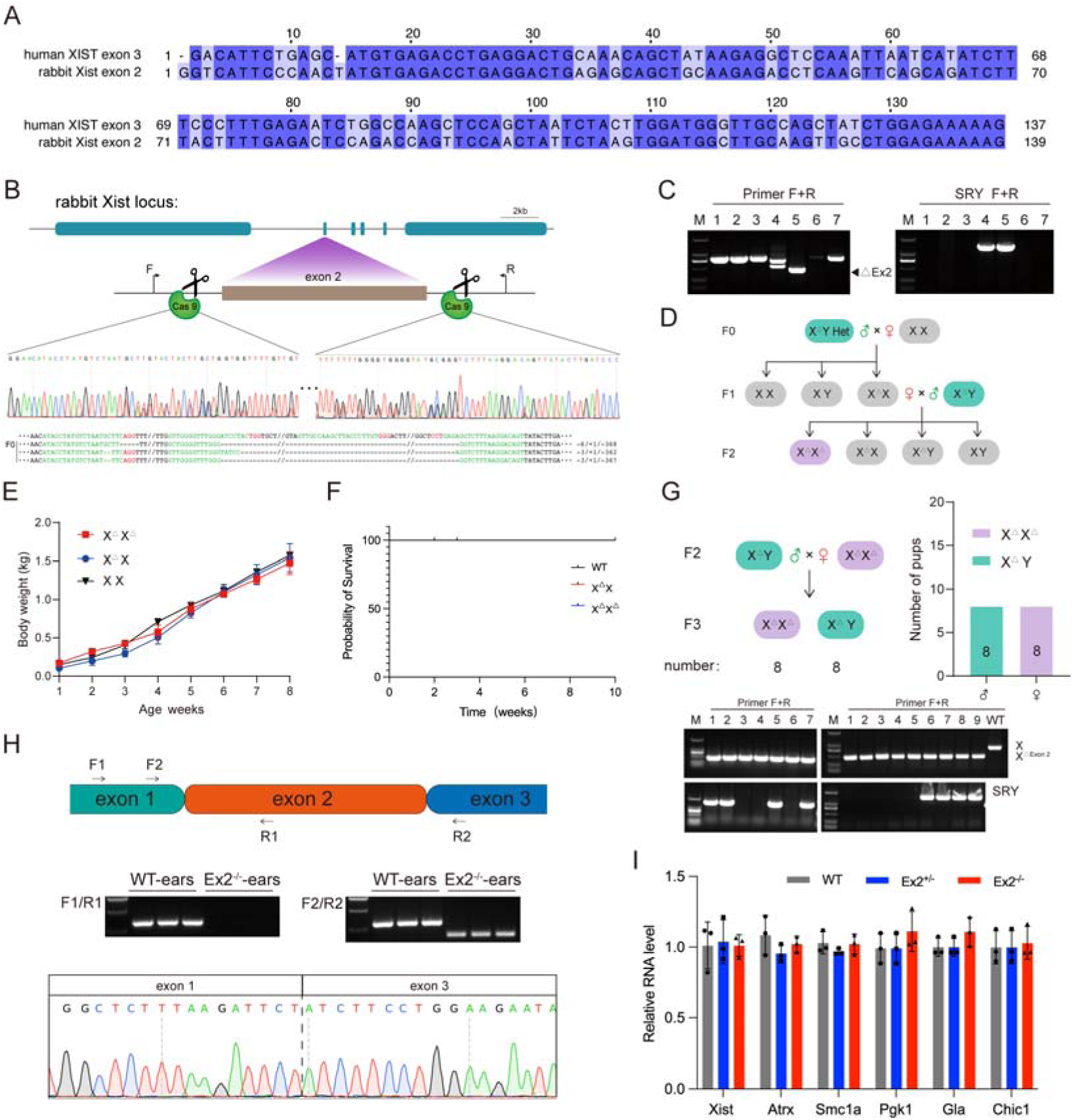
Viability of *Xist* exon 2 knockout rabbits. (**A**) Sequence alignment comparing human *XIST* exon 3 and rabbit *Xist* exon 2, with identical bases highlighted on a dark background. (**B**) Sanger sequencing results and target loci confirm *Xist* exon 2 knock-outs in F0 rabbits. All sgRNA sequences are listed in Appendix Table 1. (**C**) Agarose gel electrophoresis result of PCR product in the F0 generation. (**D**) Schematic illustrating the generation of the F1 and F2 generations. (**E**) Comparison of body weight between KO and WT rabbits, with error bars representing the mean ± SEM. (**F**) Survival curve for KO and WT rabbits. (**G, up**) Schematic diagram outlining the breeding strategy employed to generate the F3 generation. Genotype data for F3, along with the number of pups for each genotype, is provided. (**Down**) Agarose gel electrophoresis result of PCR product in the F3 generation. (**H, up**) The *Xist* expression was analyzed using RT-PCR. The agarose gel electrophoresis of the PCR products is presented. (**Down**) The Sanger sequencing results confirmed the findings from the RT-PCR analysis. (**I**) The qPCR results of X-linked genes in Ex2^+/-^ and Ex2^−/−^ rabbits are displayed. The error bars represent the mean ± SEM.

**Figure EV4.**
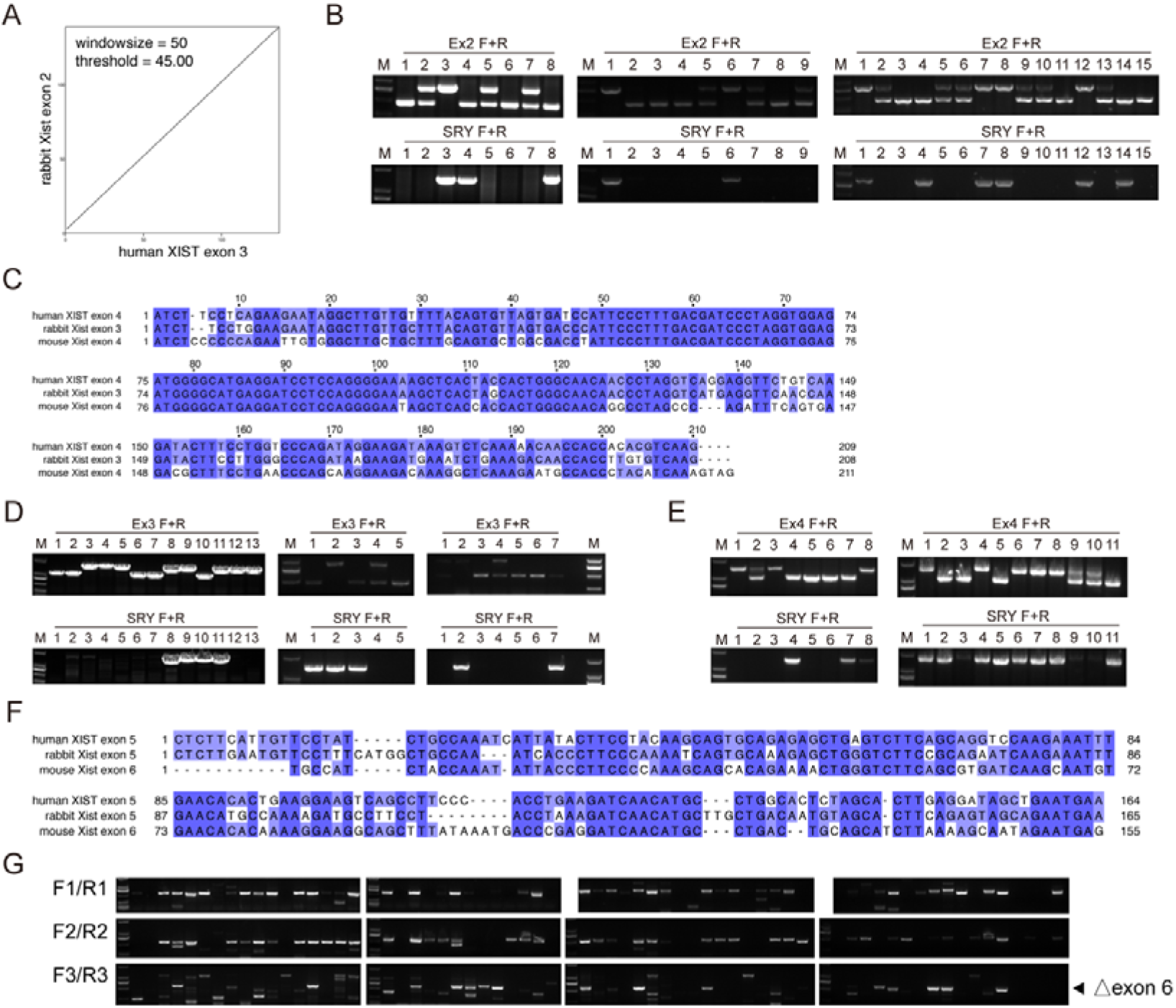
Sequence alignment and agarose gel electrophoresis results. (**A**) Dot plot analysis comparing rabbit *Xist* exon 2 and human *XIST* exon 3. (**B**) Agarose gel electrophoresis result of the PCR product (Ex2 F1 generation). (**C**) Sequence alignment of human *XIST* exon 4, rabbit *Xist* exon 3, and mouse *Xist* exon 4. Identical bases are shown on a dark background. (**D**) Agarose gel electrophoresis result of the PCR product (Ex3 F1 generation). (**E**) Agarose gel electrophoresis result of the PCR product (Ex4 F1 generation). (**F**) Sequence alignment of human *XIST* exon 5, rabbit *Xist* exon 5, and mouse *Xist* exon 6. Identical bases are shown on a dark background. (**G**) Identification of cloned embryo by agarose gel electrophoresis using primers F1/R1, F2/R2, and F3/R3.

Seven founder (F0) pups were identified through Sanger sequencing and a chimeric male with exon 2 deletion was obtained (Fig 4B and C). Through subsequent backcrossing, we successfully obtained female homozygous knockout rabbits (Ex2^−/−^) in the F2 generation (Figs 4D and EV4B). And the RT-PCR results demonstrated the absence of exon 2 sequence in the expressed *Xist* RNA of the rabbits (Fig 4H). Importantly, there were no significant differences observed in terms of body weight (Fig 4E), survival rates (Fig 4F), reproductive efficiency (Fig 4G) and X-linked gene expression (Fig 4I) between the Ex2^−/−^ and WT rabbits. These findings indicate normal growth and development of the Ex2^−/−^ rabbits.

### Deletions of *Xist* exon 3 in rabbits are viable and develop normally

Sequence homology analysis results revealed a sequence homology of rabbit *Xist* exon 3 vs. human *XIST* exon 4 was 79.80% and a coverage of 94%, which was significantly higher than the 76.80% and 89% observed in human exon 4 vs. mouse *Xist* exon 4 (Figs 5A and EV4C). These findings were also confirmed by Dot plot analysis (Fig 5B). Thus, the deletions of *Xist* exon 3 in rabbits were used to mimic the function of *XIST* exon 4 in humans.

**Figure 5.**
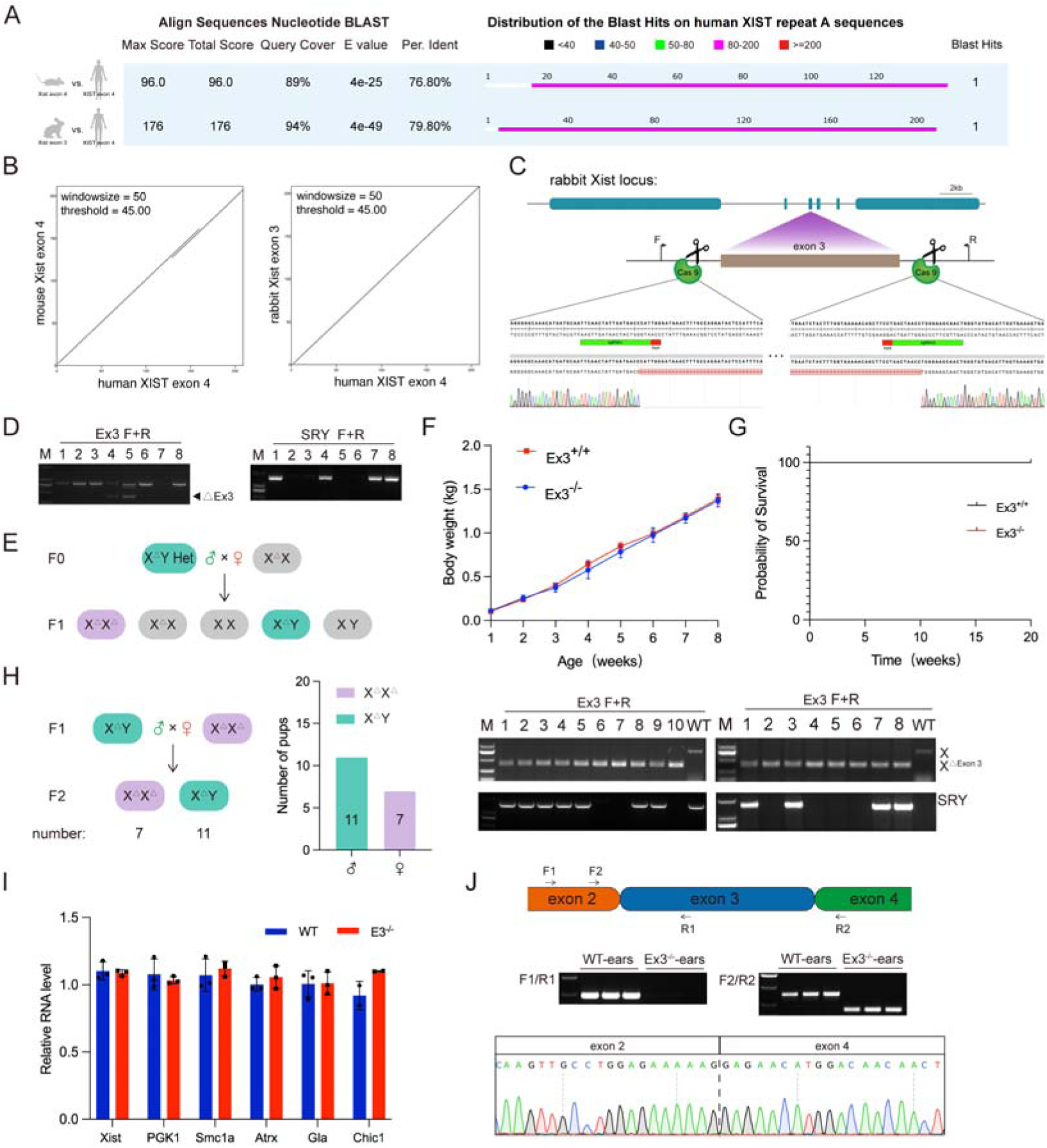
Viability of *Xist* exon 3 knockout rabbits. (**A**) BLAST results comparing human *XIST* exon 4, rabbit *Xist* exon 3, and mouse *Xist* exon 4. (**B**) Dot plot analysis of mouse, rabbit, and human sequences. (**C**) Sanger sequencing results and target loci confirm *Xist* exon 3 knock-outs in F0 rabbits. All sgRNA sequences are listed in Appendix Table 1. (**D**) Agarose gel electrophoresis shows PCR product of F0 generation. (**E**) Schematic representation of generating the F1 generation. (**F**) Comparison of body weight between KO and WT rabbits. Error bars represent mean ± SEM. (**G**) Survival curve comparing KO and WT rabbits. (**H, left**) Schematic diagram illustrating the breeding strategy employed to generate the F2 generation. Genotype data for the F2 generation is provided, listing the number of pups for each genotype. (**Right**) Result of agarose gel electrophoresis showing the PCR product for the F2 generation. (**I**) qPCR result displaying the expression of X-linked genes in Ex3^−/−^ rabbits. Error bars represent the mean ± SEM. (**J, up**) RT-PCR analysis of *Xist* expression, with agarose gel electrophoresis of the PCR products shown. (**Down**) The Sanger sequencing results support the RT-PCR analysis findings.

The founder (F0) pups were identified through Sanger sequencing and a male chimeric rabbit with exon 3 knockout and a single knockout female rabbit was obtained (Fig 5C and D). Through subsequent breeding, we successfully obtained female homozygous knockout rabbits (Ex3^−/−^) in the F1 generation (Figs 5E and EV4D). And the RT-PCR results demonstrated the absence of exon 3 sequence in the expressed *Xist* RNA of the rabbits (Fig 5J). Importantly, there were no significant differences observed in terms of body weight (Fig 5F), survival rates (Fig 5G), reproductive efficiency (Fig 5H) and X-linked gene expression (Fig 5I) between the Ex3^−/−^ and WT rabbits. These findings indicate normal growth and development of the Ex3^−/−^ rabbits.

### Deletions of *Xist* exon 4 in rabbits are viable and develop normally

Rabbit *Xist* exon 4 is the only exon that does not have sequence homology with its human counterpart. In order to investigate the functionality of rabbit *Xist* exon 4, we used a targeted approach involving a pair of sgRNAs to specifically disrupt this exon. The F0 generation rabbits were identified using Sanger sequencing and a male chimeric individual with a complete knockout of exon 4 and a single knockout female individual were obtained (Fig 6A and B). Through subsequent breeding, we successfully obtained female homozygous knockout rabbits (Ex4^−/−^) in the F1 generation (Figs 6C and EV4E). And the RT-PCR results demonstrated the absence of exon 4 sequence in the expressed *Xist* RNA of the rabbits (Fig 6G). Importantly, there were no significant differences observed in terms of body weight (Fig 6D), survival rates (Fig 6E), reproductive efficiency (Fig 6F) and X-linked gene expression (Fig 6H) between the Ex4^−/−^ and WT rabbits. These findings indicate normal growth and development of the Ex4^−/−^ rabbits.

**Figure 6.**
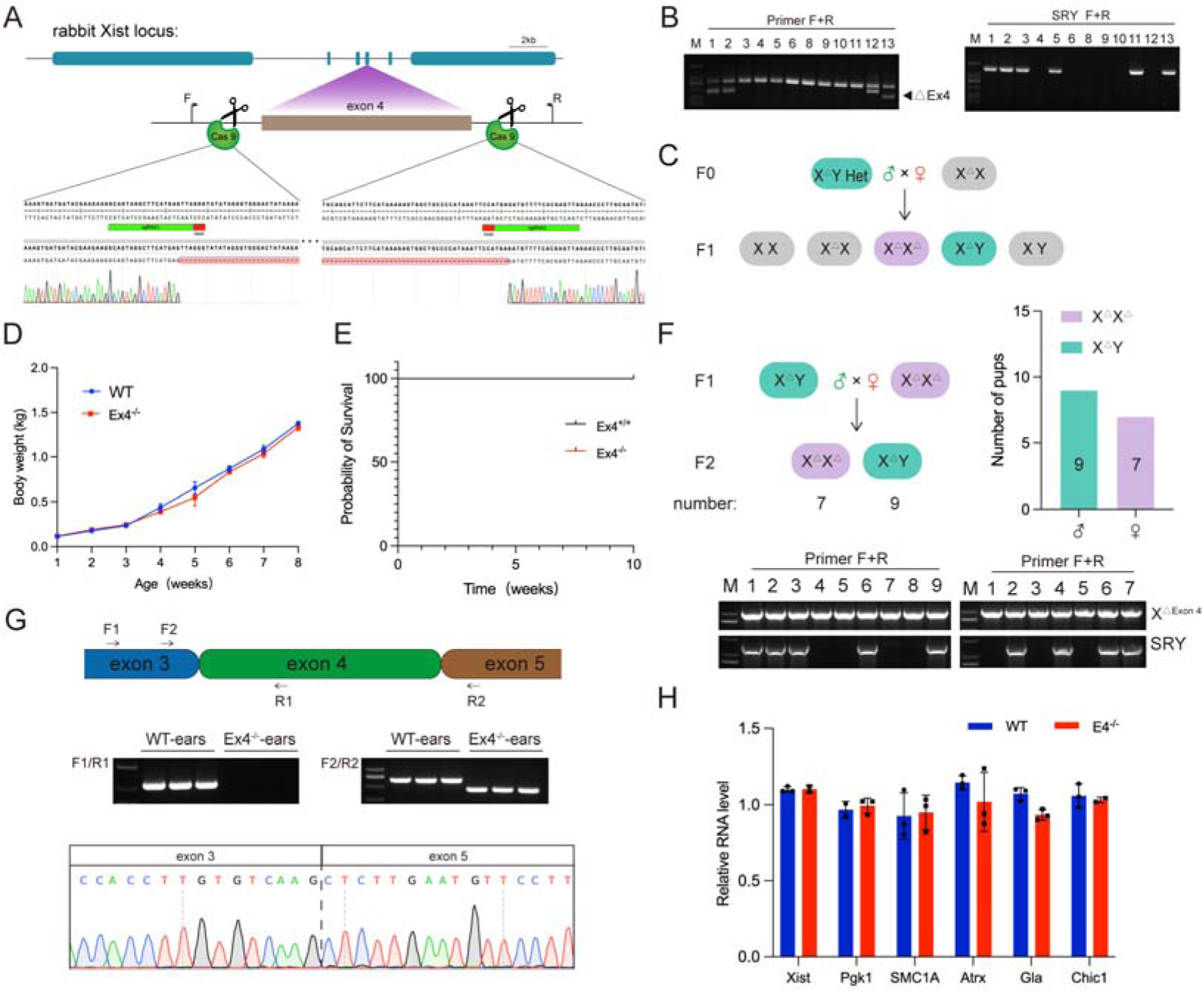
Viability of *Xist* exon 4 knockout rabbits. (**A**) Sanger sequencing results and target loci confirm *Xist* exon 4 knockouts in F0 rabbits. All sgRNA sequences are listed in Appendix Table 1. (**B**) Agarose gel electrophoresis shows the PCR product of the F0 generation. (**C**) Schematic representation of the generation process of the F1 generation. (**D**) Body weight comparison between KO and WT rabbits. Error bars represent mean ± SEM. (**E**) Survival curve for KO and WT rabbits. (**F, up**) Breeding strategy schematic for generating the F2 generation. Genotype data and number of pups for each genotype are provided. (**Down**) Agarose gel electrophoresis result of the PCR product from the F2 generation. (**G, up**) RT-PCR analysis of *Xist* expression. Agarose gel electrophoresis shows the PCR products. (**Down**) The RTPCR analysis results are supported by Sanger sequencing. (**H**) The qPCR result of X-linked genes in Ex4^−/−^ rabbits. Error bars represent mean ± SEM.

### Deletions of *Xist* exon 5 in rabbits are viable and develop normally

Sequence homology analysis results revealed a sequence homology of rabbit *Xist* exon 5 vs. human *XIST* exon 5 was 74.42% and a coverage of 100%, which was significantly higher than the 74.59% and 78% observed in human exon 5 vs. mouse *Xist* exon 6 (Figs 7A and EV4F). These findings were also confirmed by Dot plot analysis (Fig 7B). Thus, the deletions of *Xist* exon 5 in rabbits were used to mimic the function of *XIST* exon 5 in humans.

**Figure 7.**
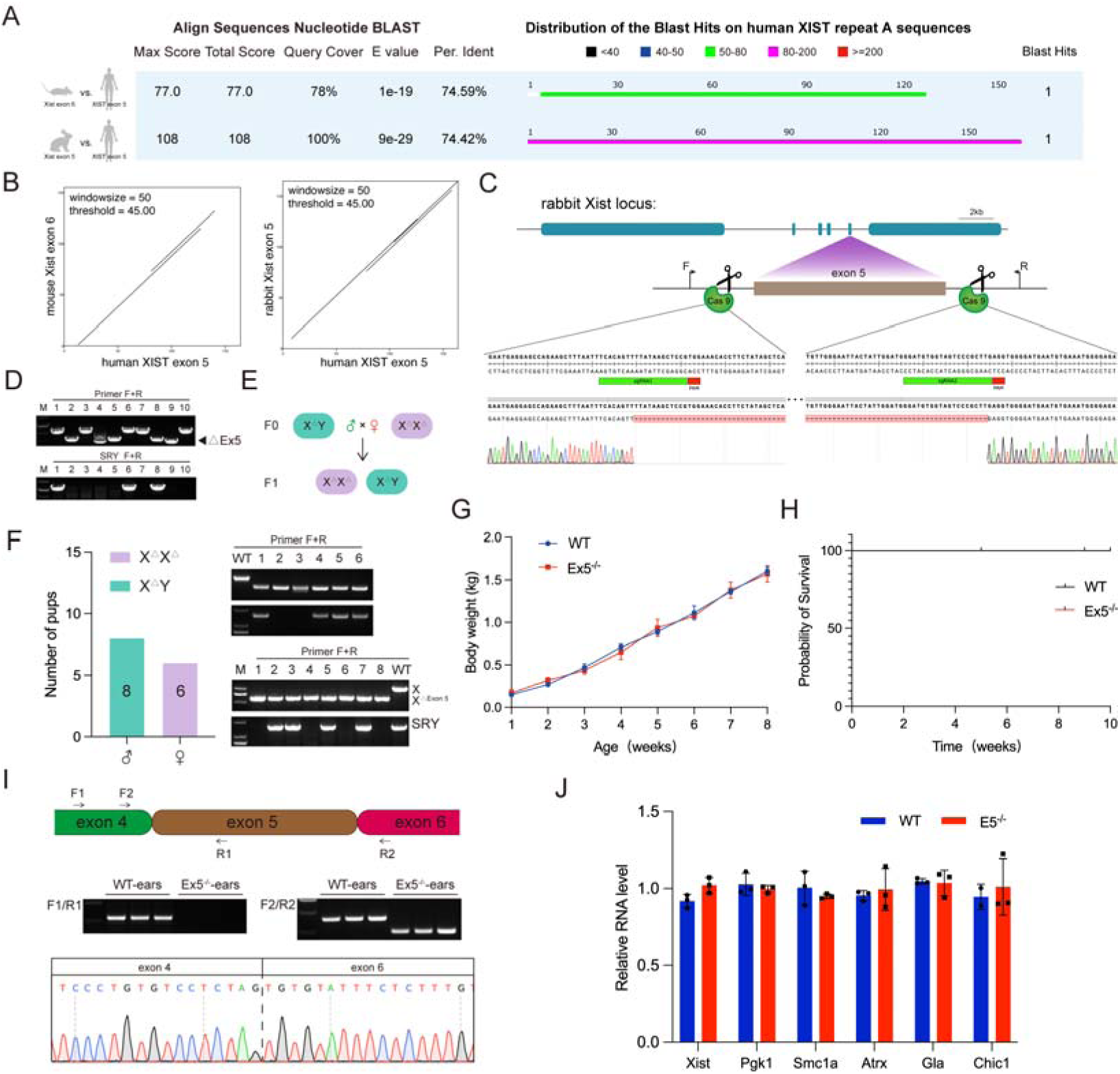
Viability of *Xist* exon 5 knockout rabbits. (**A**) BLAST analysis comparing human *XIST* exon 5, rabbit *Xist* exon 5, and mouse *Xist* exon 6. (**B**) Dot plot analysis of mouse, rabbit, and human sequences. (**C**) Sanger sequencing results and target loci confirm *Xist* exon 5 knock-outs in F0 rabbits. All sgRNA sequences are listed in Appendix Table 1. (**D**) Agarose gel electrophoresis result of PCR product in the F0 generation. (**E**) Schematic representation of the process for generating the F1 generation. (**F, left**) Genotype data for F2 rabbits; the number of pups for each genotype is provided. (**Right**) Agarose gel electrophoresis result of PCR product in the F2 generation. (**G**) The body weight of KO and WT rabbits. Error bars represent mean ± SEM. (**H**) The survival curve for KO and WT rabbits. (**I, up**) RT-PCR analysis of the *Xist* expression. Agarose gel electrophoresis of the PCR products is shown. (**Down)** The Sanger sequencing results supported the RTPCR analysis results. (**J**) qPCR result of X-linked genes in Ex5^−/−^ rabbits. Error bars represent mean ± SEM.

The founder (F0) pups were identified through Sanger sequencing and homozygous exon 5 knockout male and female rabbits were obtained (Fig 7C and D). Subsequent breeding led to the generation of F1 rabbits (Fig 7E). And the RT-PCR results demonstrated the absence of exon 5 sequence in the expressed *Xist* RNA of the rabbits (Fig 7I). Importantly, there were no significant differences observed in terms of reproductive efficiency (Fig 7F), body weight (Fig 7G), survival rates (Fig 7H) and X-linked gene expression (Fig 7J) between the Ex5^−/−^ and WT rabbits. These findings indicate normal growth and development of the Ex5^−/−^ rabbits.

### Deletion of *Xist* exon 6 in rabbits results in embryonic lethality

The results of the sequence homology analysis showed that there was a 70.13% sequence homology between rabbit *Xist* exon 6 and human *XIST* exon 6, with a coverage of 95%. This was significantly higher compared to the 70.85% sequence homology and 39% coverage observed between human exon 6 and mouse *Xist* exon 7 (Fig 8A). These findings were also confirmed by Dot plot analysis (Fig 8B). Thus, the deletions of *Xist* exon 6 in rabbits were used to mimic the function of *XIST* exon 6 in humans.

**Figure 8.**
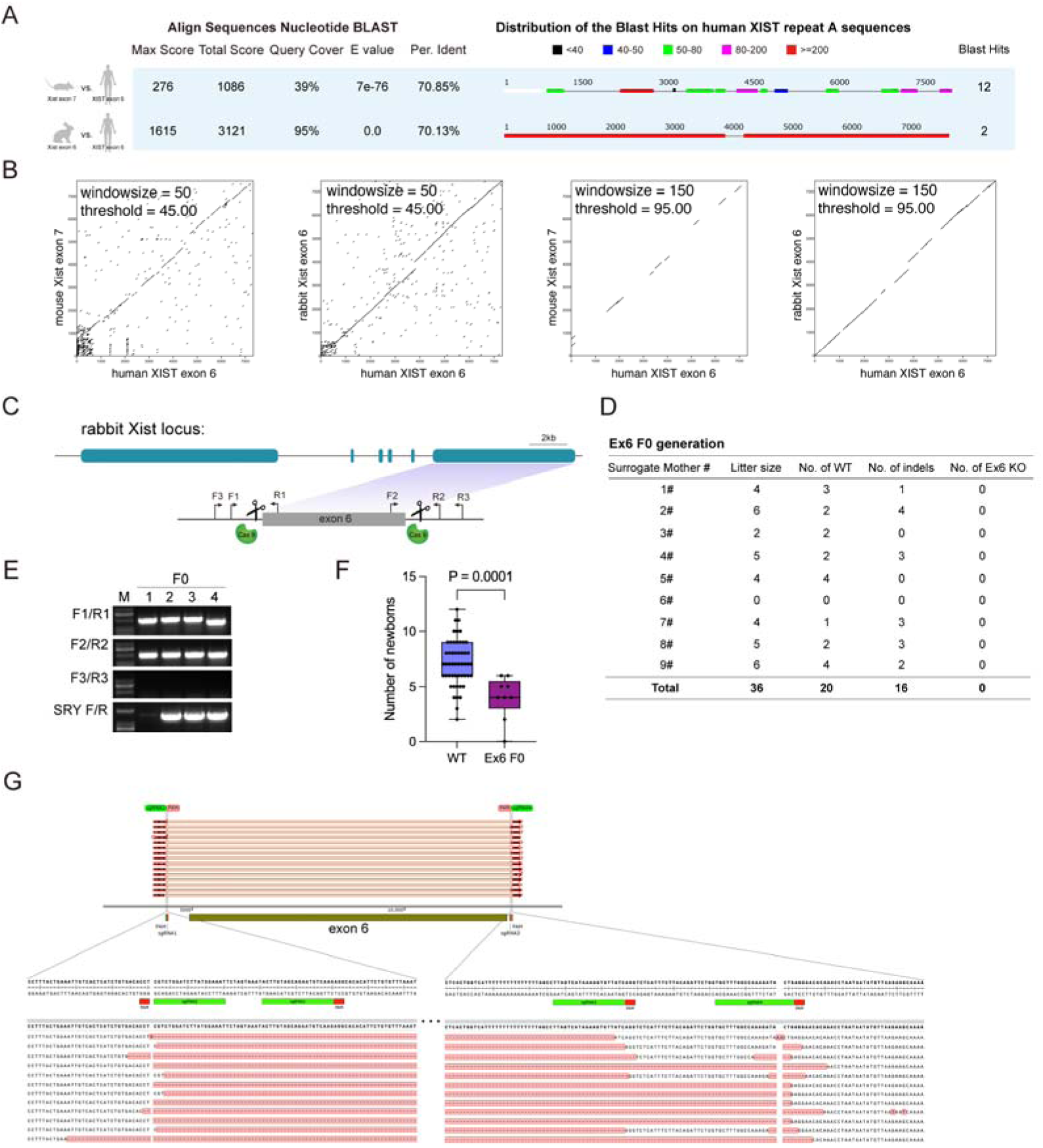
*Xist* exon 6 knockout rabbits do not survive. (**A**) The BLAST result shows the comparison of human *XIST* exon 6, rabbit *Xist* exon 6, and mouse *Xist* exon 7. (**B**) Dot plot analysis illustrates the sequences of mouse, rabbit, and human. (**C**) The map indicates the target sites of CRISPR/Cas9 in the *Xist* loci. All sgRNA sequences are listed in Appendix Table 1. (**D**) A table summarizes the litter size of live-born offspring in different genotypes of the F0 generation. (**E**) Representative images of agarose gel electrophoresis display the PCR products from the F0 generation. (**F**) The newborn data from the F0 generation is compared to the WT control using a box and whiskers plot, showing the minimum to maximum values and t-test results (P=0.0001). (**G**) The target loci and Sanger sequencing results confirm the knock-out of *Xist* exon 6 in rabbit embryos.

To disrupt the function of *Xist* exon 6 in rabbits, we designed two pairs of sgRNAs targeting *Xist* exon 6 (Fig 8C). The F0 generation rabbits were subsequently confirmed through PCR and Sanger sequencing. Surprisingly, no *Xist* exon 6 knockout animals were obtained from pregnant females (Fig 8D and E), indicating that the absence of exon 6 resulted in non-viability. Additionally, statistical analysis of offspring production revealed a reduced number of offspring from pregnant females compared to WT rabbits (Fig 8F).

To further characterize the lethal stage of early embryonic development, we injected Cas9 mRNA and sgRNA into fertilized zygotes. Surprisingly, we were able to successfully knockout rabbit *Xist* exon 6 in early-stage embryos (Figs 8G and EV4G). However, our results revealed that the cloned embryos had a significantly lower blastocyst rate (13.3±0.9%) compared to the control group (74.7±5.4%). These findings provide conclusive evidence that the deletion of *Xist* exon 6 leads to embryonic lethality in rabbits, hindering proper development.

## Discussion

*Xist* is continuously expressed in female somatic cells to maintain the state of X chromosome inactivation (XCI) (Yang *et al*, 2022). Previous studies investigating the major functional domains of *Xist* have primarily relied on mouse models and cellular-level experiments. However, it is important to note that mice displays paternal Xist expression during the initial stage of X chromosome inactivation, followed by random X chromosome inactivation during the subsequent stage, which differs significantly from the random XCI mechanism observed in humans (Balaton *et al*, 2021; Goto & Monk, 1998b). Therefore, the consistency of cellular-level results in vivo remains uncertain. At the cellular level, numerous studies have suggested that repeat A recruits chromatin silencing factors to establish gene silencing on the X chromosome. Deletion of repeat A has been found to result in X chromosome silencing failure (Trotman *et al*, 2020; Colognori *et al*, 2020a; Coker *et al*, 2020; Sakata *et al*, 2017; Brockdorff, 2017). However, the implications of this finding in vivo remain unknown. In our study, we conducted a deletion of repeat A in rabbits and observed that individuals who were homozygous for the deletion exhibited embryonic lethality and were unable to develop. Interestingly, individuals who were heterozygous and had the repeat A deletion transcribed the *Xist* gene from the intact X chromosome and showed no phenotypic differences compared to individuals with the wild-type genotype. When we analyzed the reproductive quantities in the offspring, we found that there was a lower number compared to the control group, suggesting that X^ΔReA^X^ΔReA^ may lead to embryonic lethality. These findings indicate that repeat A plays a crucial role in establishing X-chromosome inactivation and female development.

Deletion of the 5’ conserved region of *Xist*, which includes exon 1, in mice revealed that female mice lacking *Xist* RNA were able to develop and survive until birth (Yang *et al*, 2016). However, there was a lower frequency of female births and they had a smaller size at birth, although most organ development was normal. In our study, when exon 1 was deleted in rabbits, it led to delayed development and premature death in males. Histological analysis further revealed impaired heart development. Deletion of *Xist* exon 1 had an impact on male development and led to decreased birth rates. Moreover, females were found to be incapable of surviving, indicating the crucial role of exon 1 in the process of X-chromosome inactivation in females. Specifically, the deletion of exon 1 hindered both the development and survival of females.

The functionality of regions other than *Xist* repeat A is currently poorly understood. According to a few reports, repeat B and C play a critical role in recruiting epigenetic modifier proteins to maintain the epigenetic state of XCI (Wei *et al*, 2021), while repeat E is essential for *Xist* localization and gene silencing (Cherney *et al*, 2023). However, the functions of other regions are still unknown. In a previous study, it was suggested that *XIST* exon 5 is crucial for maintaining XCI status in human K562 cells, while exons 2, 3, and 4 seem to be dispensable (Lee *et al*, 2019). However, the implications of these findings in vivo remain unknown. The results of the study indicate that rabbits lacking these exons can be born and survive normally, showing no significant differences in body weight, survival rate, or X-linked gene expression compared to WT individuals. This suggests that these exons may not be necessary for normal functioning in living organisms. Moreover, the previously emphasized importance of exon 5 in maintaining XCI status at the cellular level seems to have little significance in vivo. Additionally, the functional role of *Xist* exon 6 remains unknown. Our study found that the deletion of exon 6 in living organisms led to a lower rate of embryo blastocysts and the absence of offspring lacking exon 6. These findings demonstrate the vital and essential role of *Xist* exon 6 in embryonic development and individual survival. In conclusion, our study comprehensively elucidated the functional roles of *Xist* exons and repeat A in vivo, enhancing our understanding of the functional landscape of different regions within *Xist* and offering new insights into the functional mechanisms of *Xist* in X chromosome inactivation.

## Materials and Methods

### Animals care and use

The Institutional Animal Care and Use Committee of Jilin University approved all animal experiments. New Zealand White rabbits were obtained from the Laboratory Animal Centre of Jilin University (Changchun, China). All animal experiments were conducted by the guidelines for animal experiments of the Laboratory Animal Center of Jilin University.

### Plasmid design and construction

Eleven pairs of sgRNAs were designed to knock out *Xist* different regions according to the previous description(Liu *et al*, 2018), which were cloned into the BbsI-linearized pUC57-T7-gRNA vector. Then, sgRNAs were amplified using PCR with T7 primers (T7-Fwd: 5‘-GAA ATT AAT ACG ACT CAC TAT A-3’ and T7-Rev: 5‘-AAA AAA AGC ACC GAC TCG GTG CCA C-3’) and in vitro transcribe using the MAXIscript T7 kit (Invitrogen) and purified with a miRNeasy mini kit (QIAGEN) according to the manufacturer’s instructions. To produce SpCas9 mRNA, the PCS2+ Cas9 (Plasmid #122948) plasmid was linearized with NotI restriction digestion and used as a template to in vitro transcribe mRNAs using mMESSAGE mMACHINE SP6 Transcription Kit (Invitrogen) and then Cas9 mRNAs were purified with a miRNeasy mini kit (QIAGEN) according to the manufacturer’s instructions. All sgRNA sequences are listed in Appendix Table 1.

### Microinjection of rabbit zygotes

The protocol for sgRNA/Cas9 mRNA microinjection into pronuclear stage embryos is described in detail in our previously published study (Song *et al*, 2016). Briefly, A mixture of Cas9 mRNA (200 ng/ul) and sgRNA (50 ng/ul) was co-injected into the cytoplasm of pronuclear stage zygotes. Finally, 40-50 injected zygotes were transferred into the oviduct of recipient rabbits.

### Single-embryo and rabbit genotyping by PCR

The injected sgRNA/Cas9 mRNA zygotes were cultured for 4 days and then collected for genotyping analysis. Embryos were incubated in lysis buffer at 50℃ for 20 min and 90℃ for 5 min in a PCR machine. Genomic DNA was extracted from newborn rabbits for PCR genotyping and subjected to Sanger sequencing and T-A cloning. All primers are listed in Appendix Table 1.

### RT-PCR and Quantitative real-time PCR analysis

Tissue RNA was extracted with TRIzol (Invitrogen) according to the manufacturer’s instructions, and cDNA synthesis was performed on extracted RNA using FastKing cDNA First Strand Synthesis Kit (TIANGEN, KR116). A QuantStudio 3 Real-Time PCR System (Thermo Fisher Scientific) was used for quantitative real-time PCR experiments. Three biological replicates and three technical replicates (3 × 3) were performed for each gene. GAPDH gene was used as an internal control to normalize expression data. The RT-PCR and qRT-PCR gene-specific primers are listed in Appendix Table 2, 3, respectively.

### Hematoxylin and eosin (H&E) staining

The hematoxylin and eosin (H&E) staining was performed according to our published protocols (Yuan *et al*, 2017). Briefly, the tissues from WT and mutant rabbits were fixed in 4% paraformaldehyde for 48h, embedded in paraffin wax, and then sectioned for slides. Then, slides were stained with hematoxylin and eosin, and viewed under a Nikon ts100 microscope.

### Statistical analysis of weight and survival

To analyze survival, we conducted regular daily monitoring of the rabbits. The survival data are from 6 KO rabbits and 6 control rabbits. Body weight was recorded weekly. All data are expressed as mean ± SEM from at least three determinations in all experiments. The data were analyzed by Student’s unpaired t-test using GraphPad Prism software. p < 0.05 indicated statistical significance (∗p < 0.05, ∗∗p < 0.01, ∗∗∗p < 0.001).

### Phylogenetic tree construction

For the phylogenetic analysis of lncRNA *Xist*, all *Xist* sequences of the different species were downloaded from the NCBI database. Maximum likelihood phylogenetic trees were constructed using MEGA with 1000 bootstrap replicates (Felsenstein, 1985; Saitou & Nei, 1987; Tamura *et al*, 2004). The tree is drawn to scale, with branch lengths (next to the branches) in the same units as the evolutionary distances used to infer the phylogenetic tree (Felsenstein, 1985).

### Dot plots

To access a sequence similarity, dot plots were generated using EMBOSS dot-matcher (Madeira *et al*, 2022). Two different thresholds were used to generate the different dot plots for clarity of visualization. In this analysis, all positions of the first input sequence were systematically compared with all positions of the second input sequence using a specified substitution matrix. The resulting dot plots were generated as a rectangular grid, with the two sequences serving as the axes. Each dot in the plot represents a position where a similarity was detected between the corresponding positions of the two sequences.

### Motif analysis

Conserved motifs of each *Xist* repeat domain were defined using the MEME program (Bailey *et al*, 2015). Run with the following non-default parameters: meme sequences.fa -dna -oc. -nostatus -time 14400 -mod anr -nmotifs 100 -minw 4 -maxw 15 or 12 -objfun classic -minsites 2 maxsites 600 -revcomp -markov_order 0

### RNA secondary structure

The secondary structure graph is created from RNA secondary structure predictions using ViennaRNA Package (Lorenz *et al*, 2011). The red circles (bases) represent 90% and greater confidence with minimum free energy (MFE) and partition function.

## Acknowledgments

We thank Peiran Hu and Nannan Li for their assistance at the Embryo Engineering Center for the critical technical assistance.

## Author contributions

M.L. and Z.L. designed the experiments. M.L. and L.Z. performed the experiments. M.L., L.L., and Z.L. analyzed the data. M.L. and Z.L. wrote the paper.

---

**Table 1.**
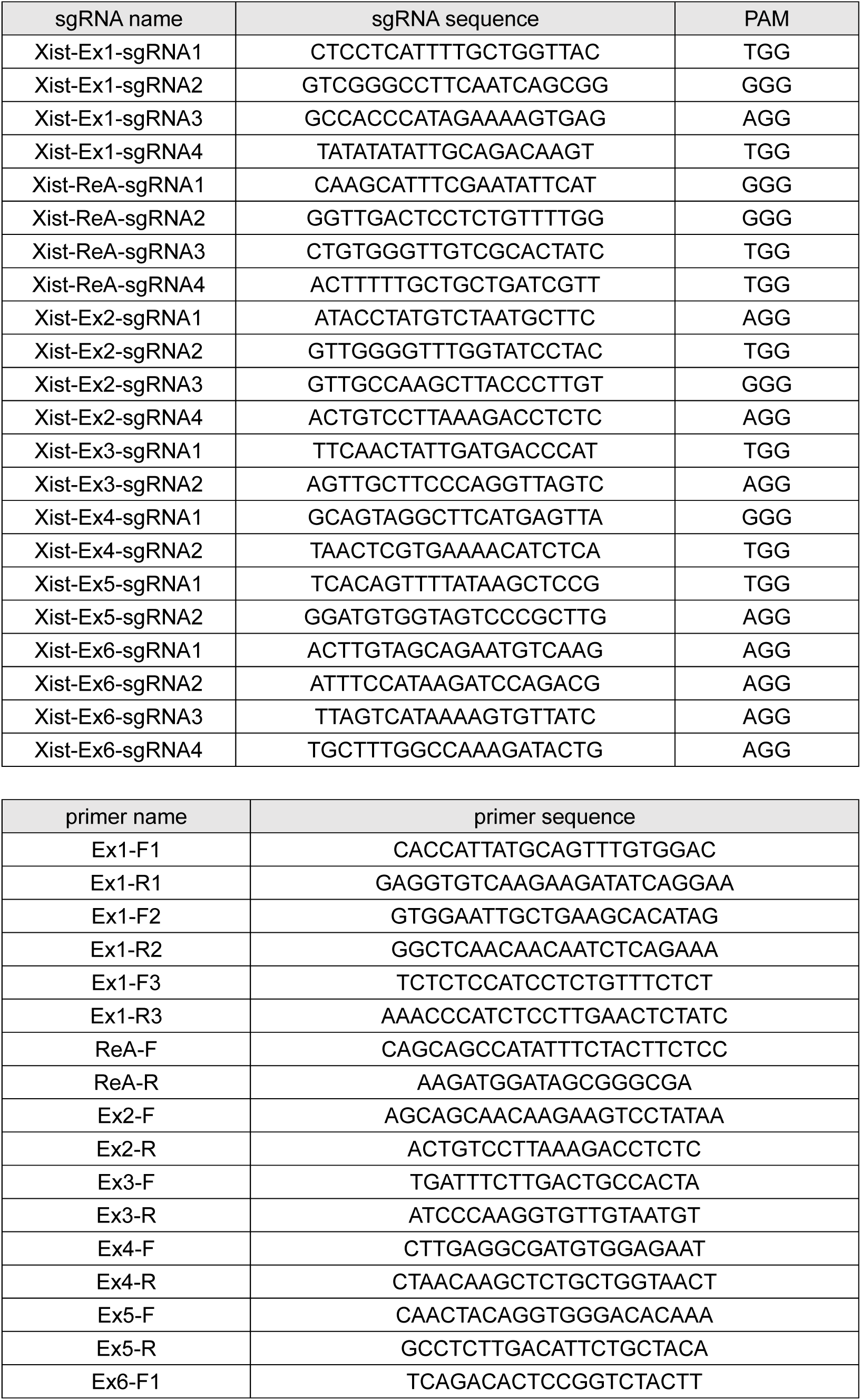
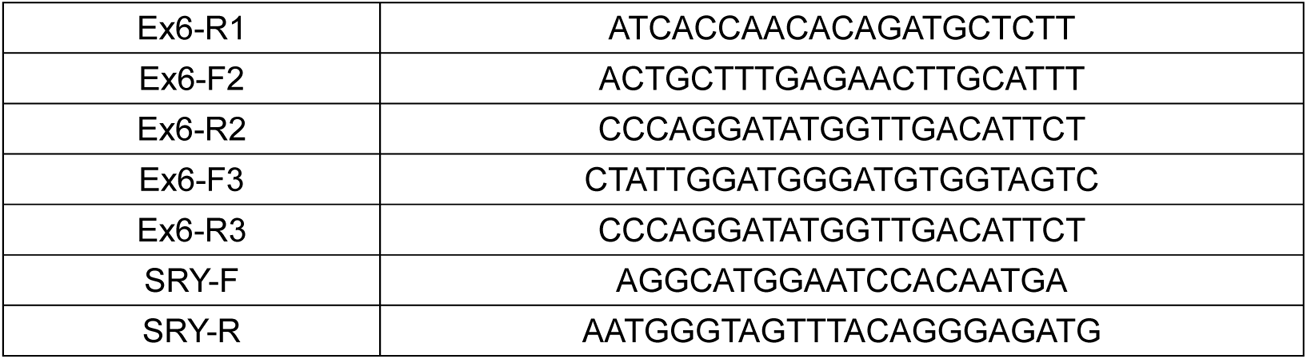
Summary of sgRNA and primer sequences used in this study.

**Table 2.**
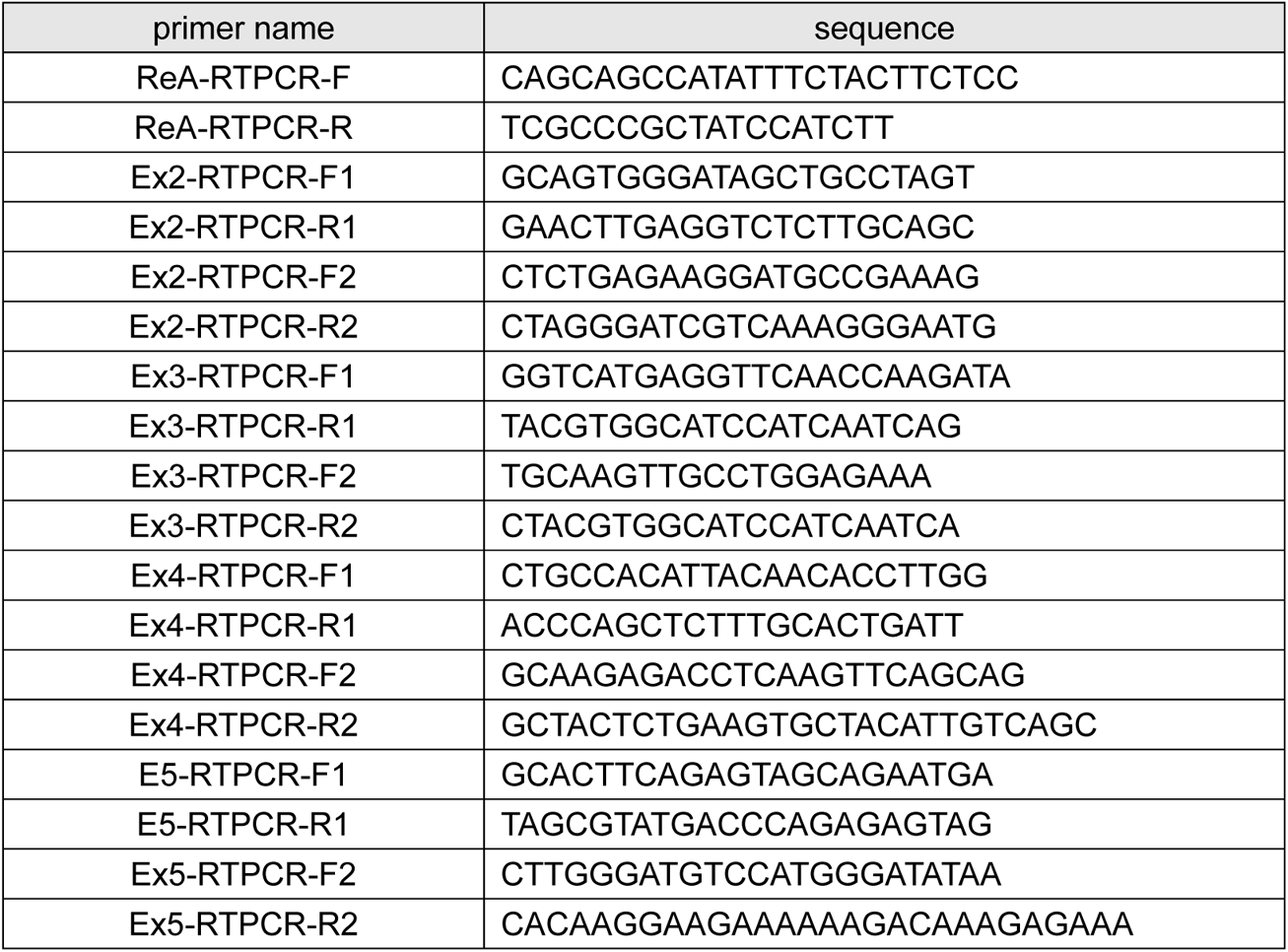
RTPCR primer sequences.

**Table 3.**
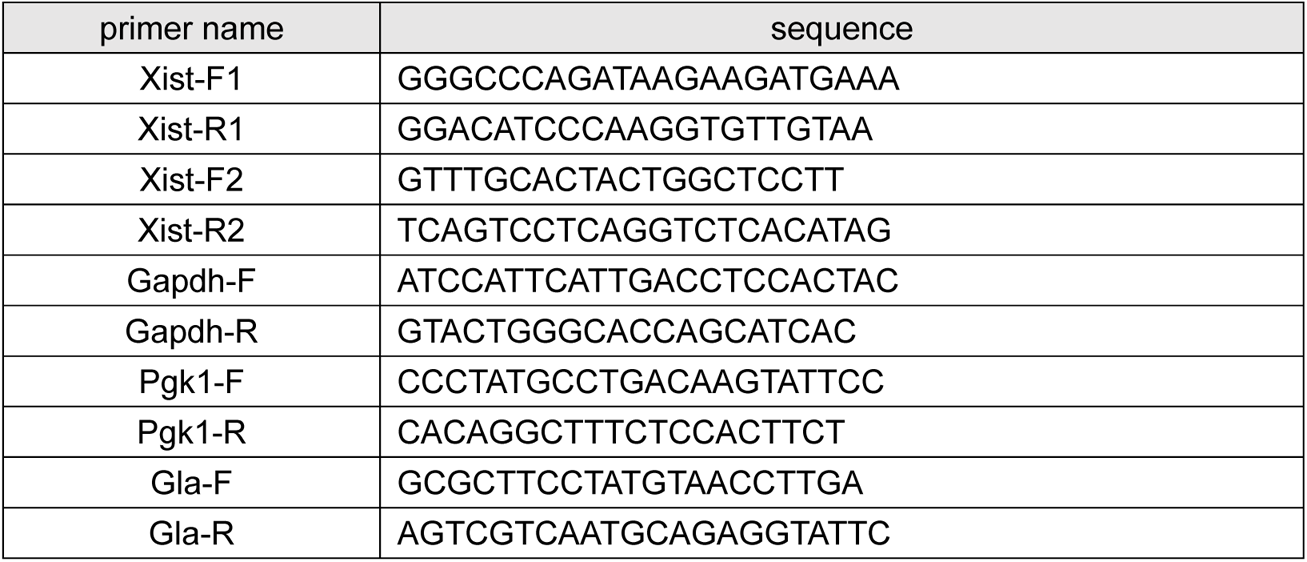
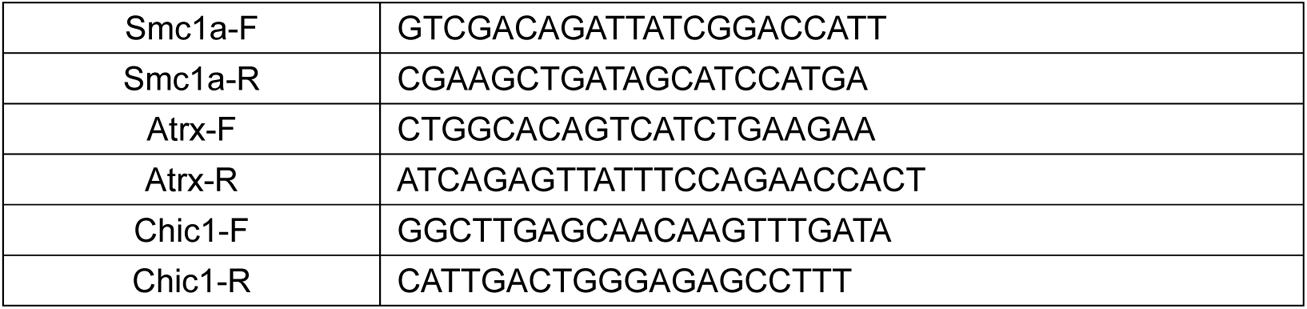
qRTPCR primer sequences.

**Table 4.**
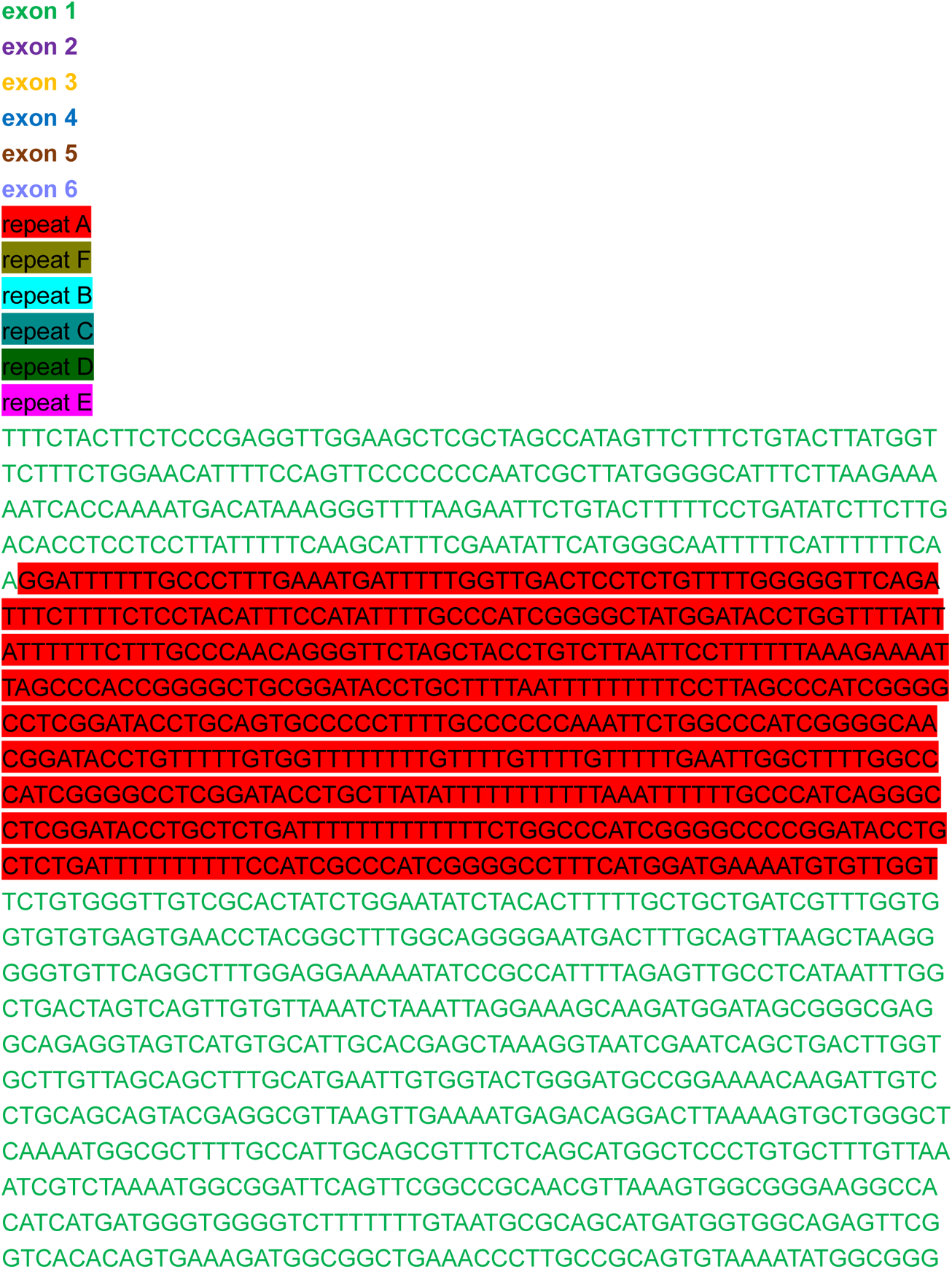
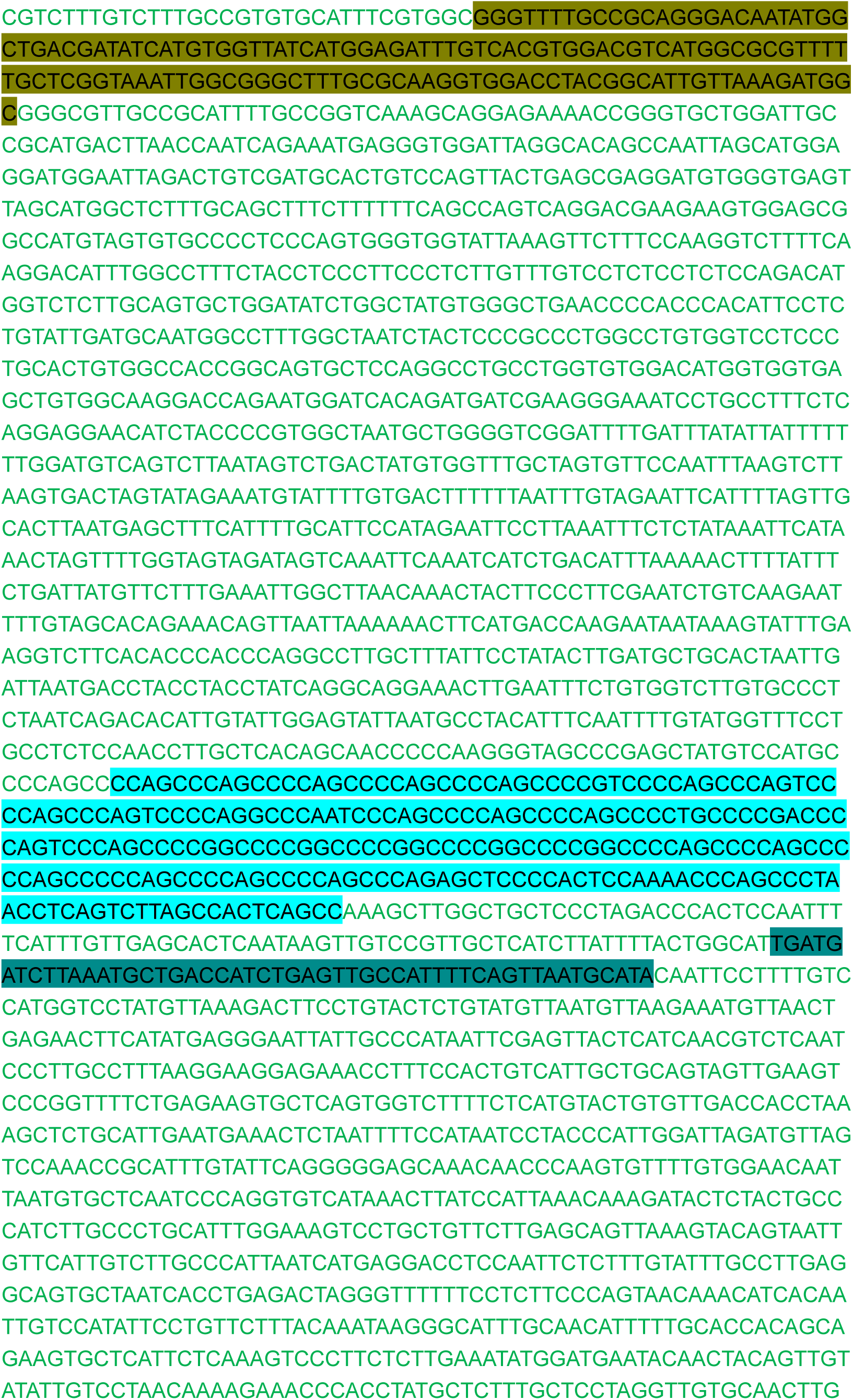
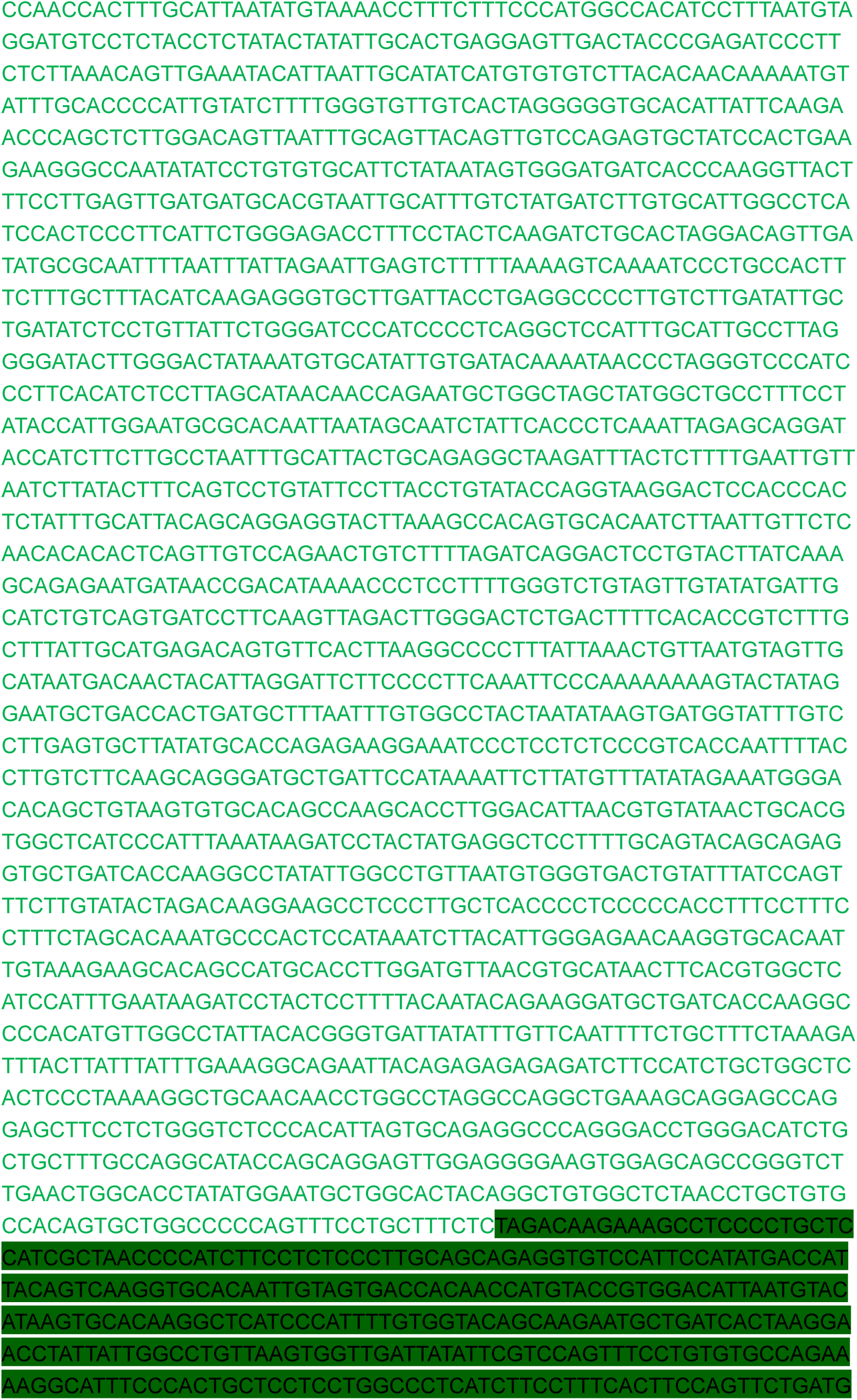
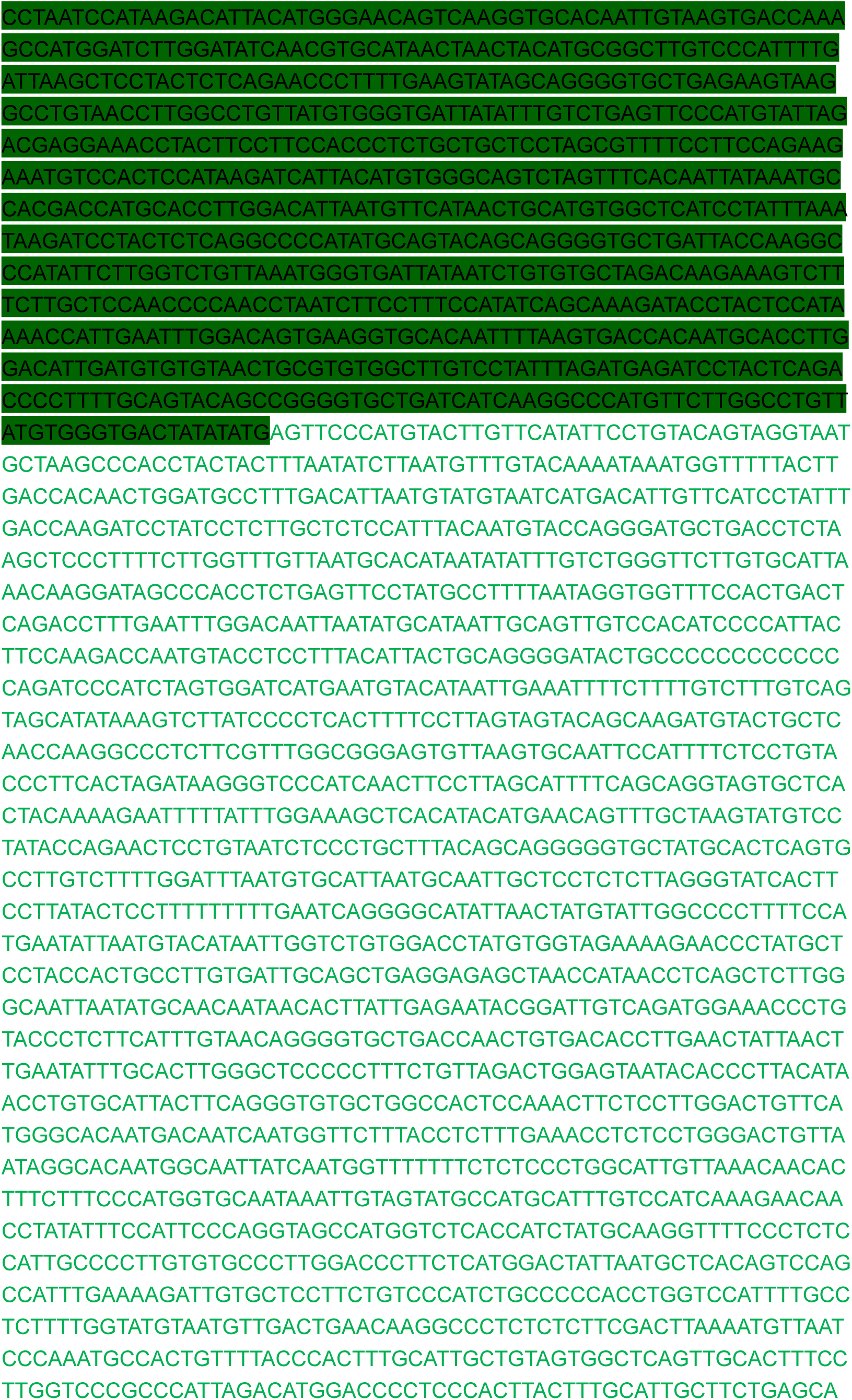
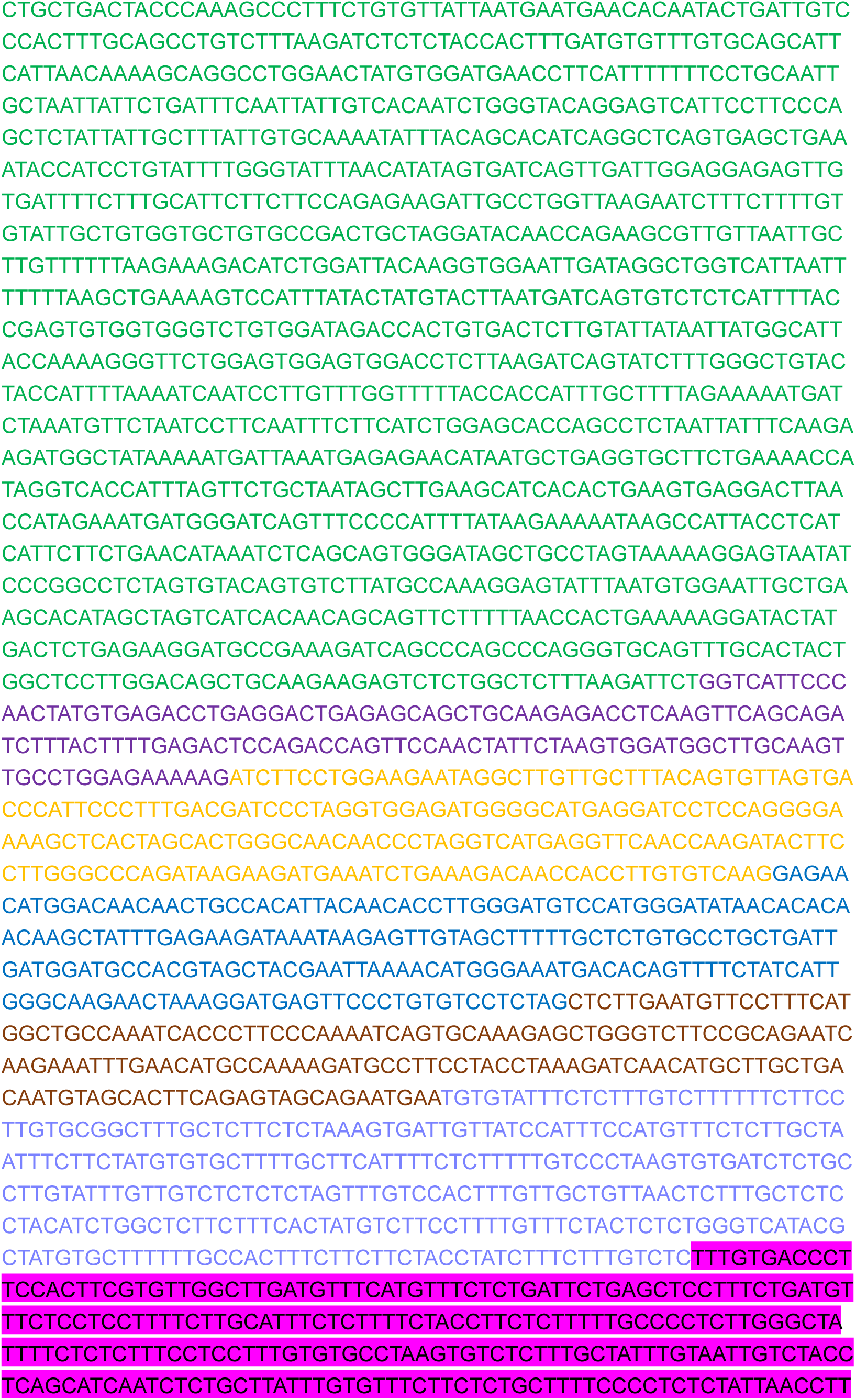
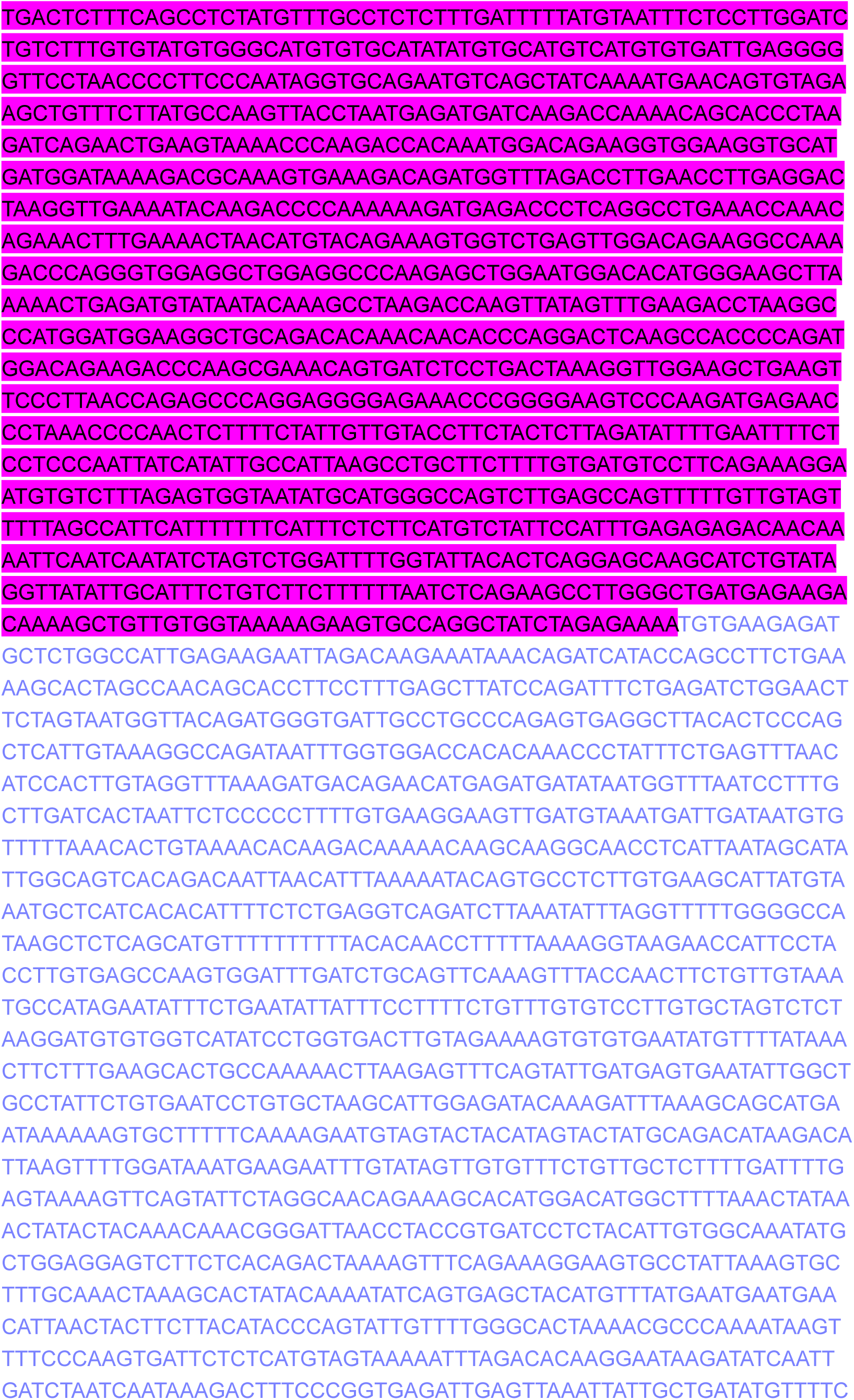
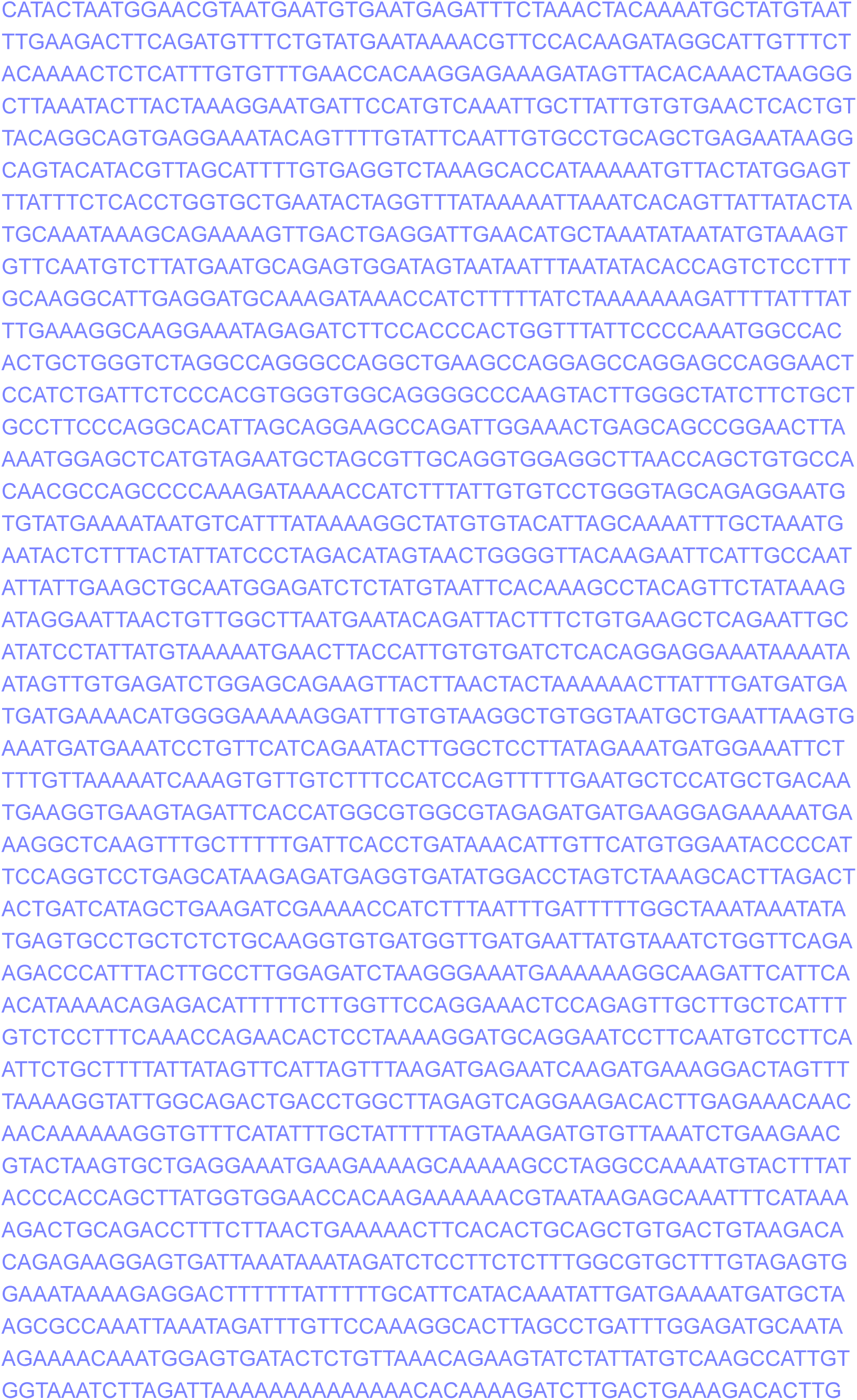
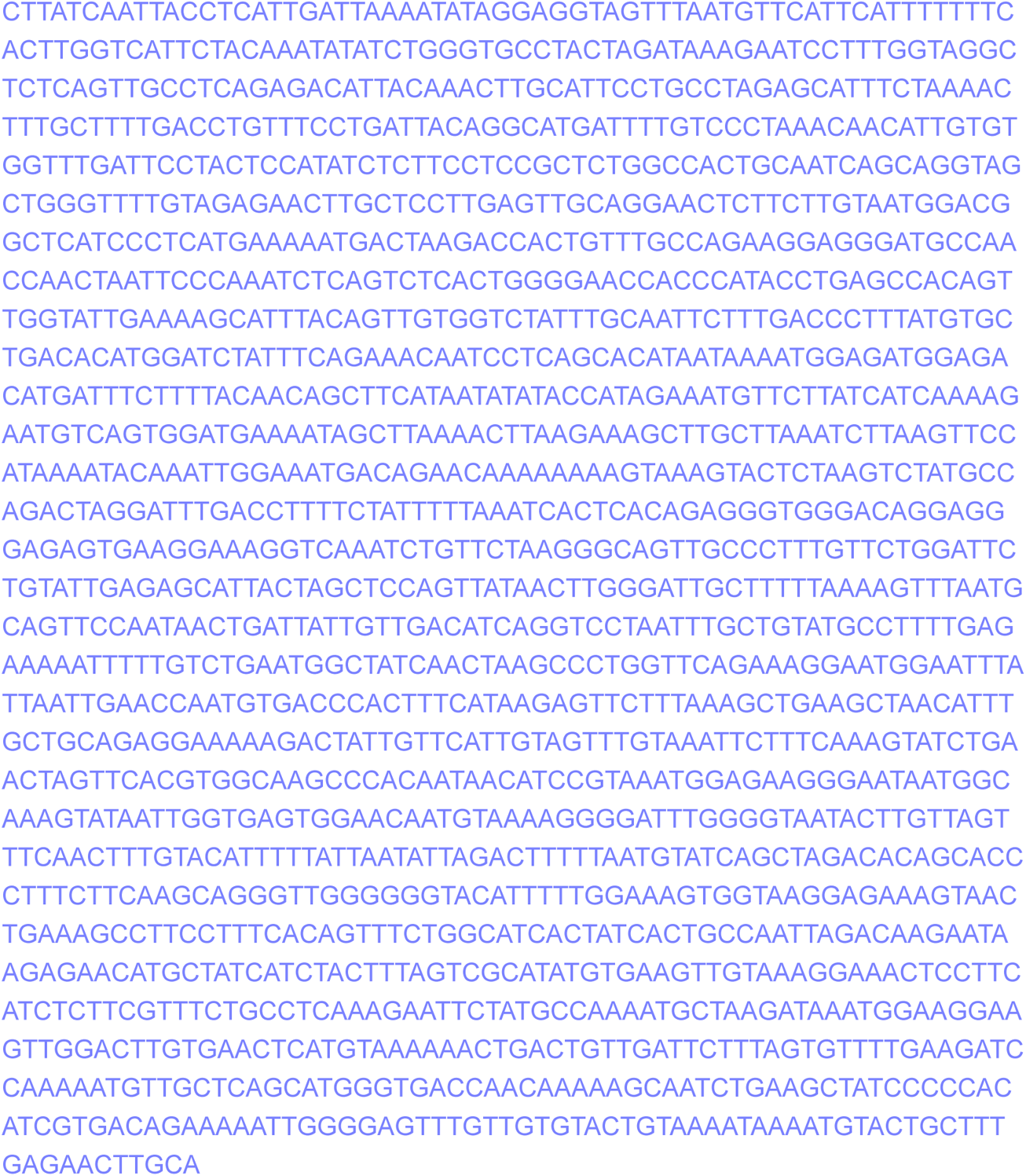
Sequence of the rabbit *Xist* locus.

**Table 5.**
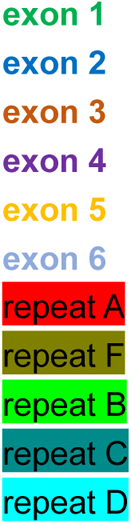
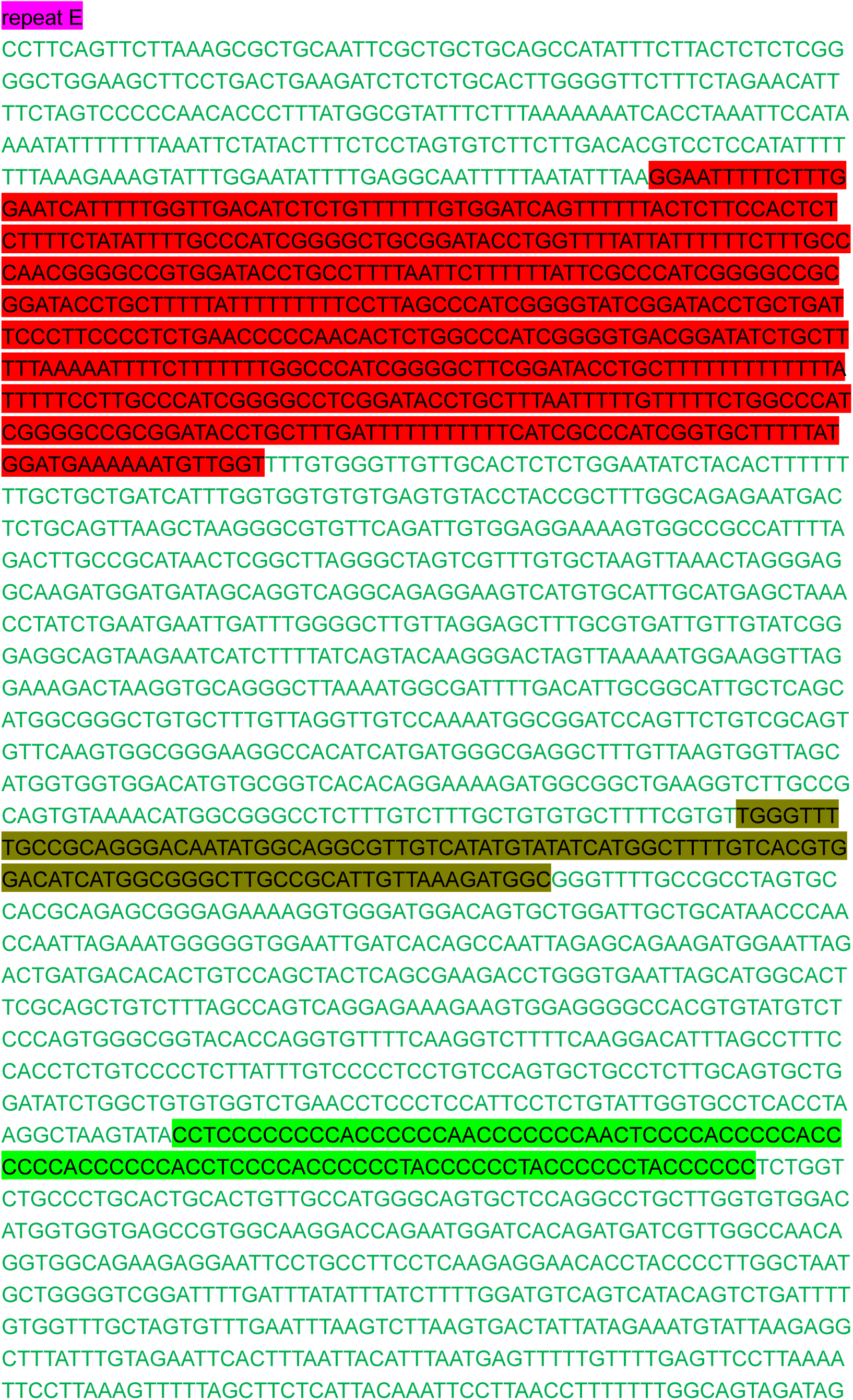
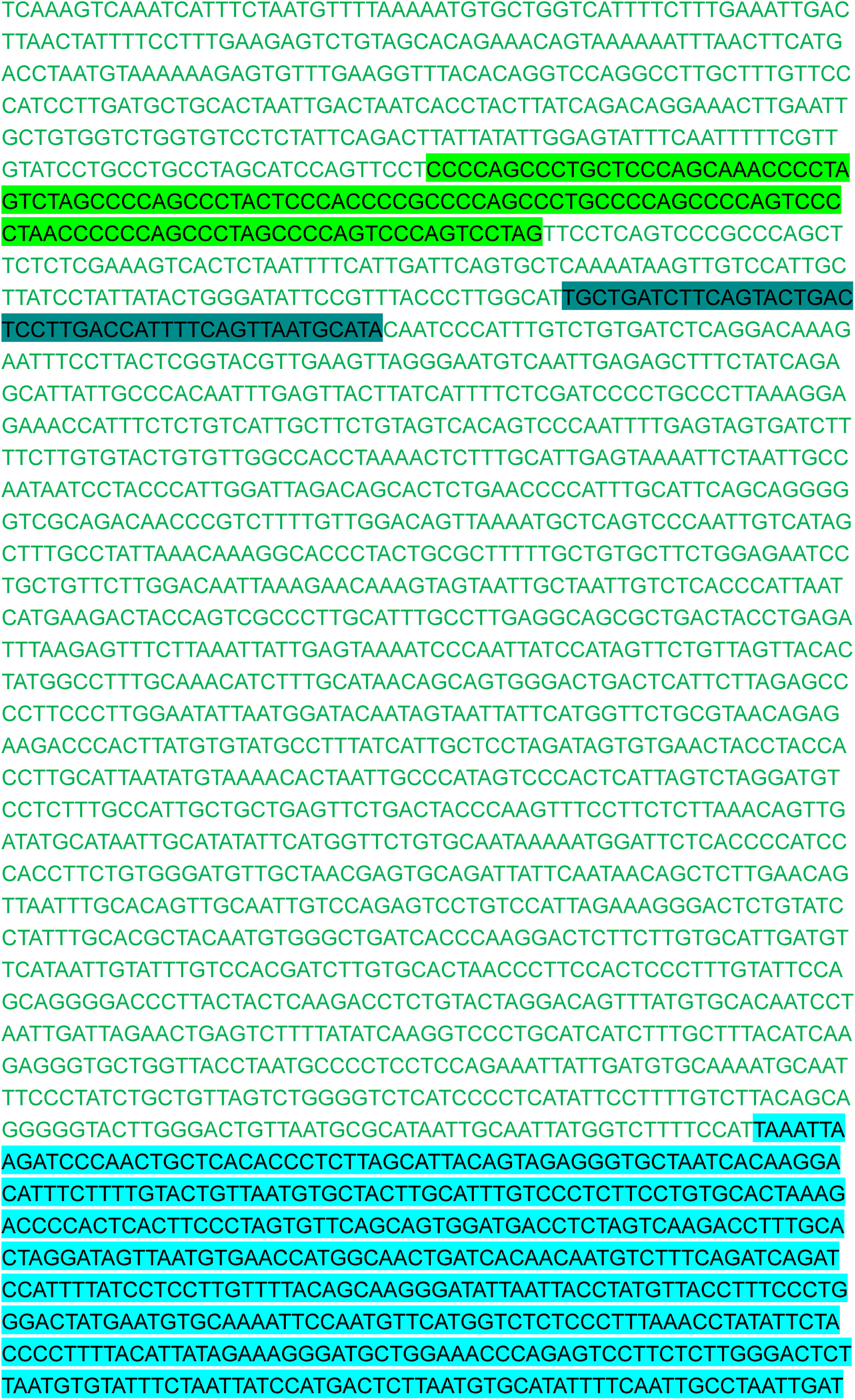
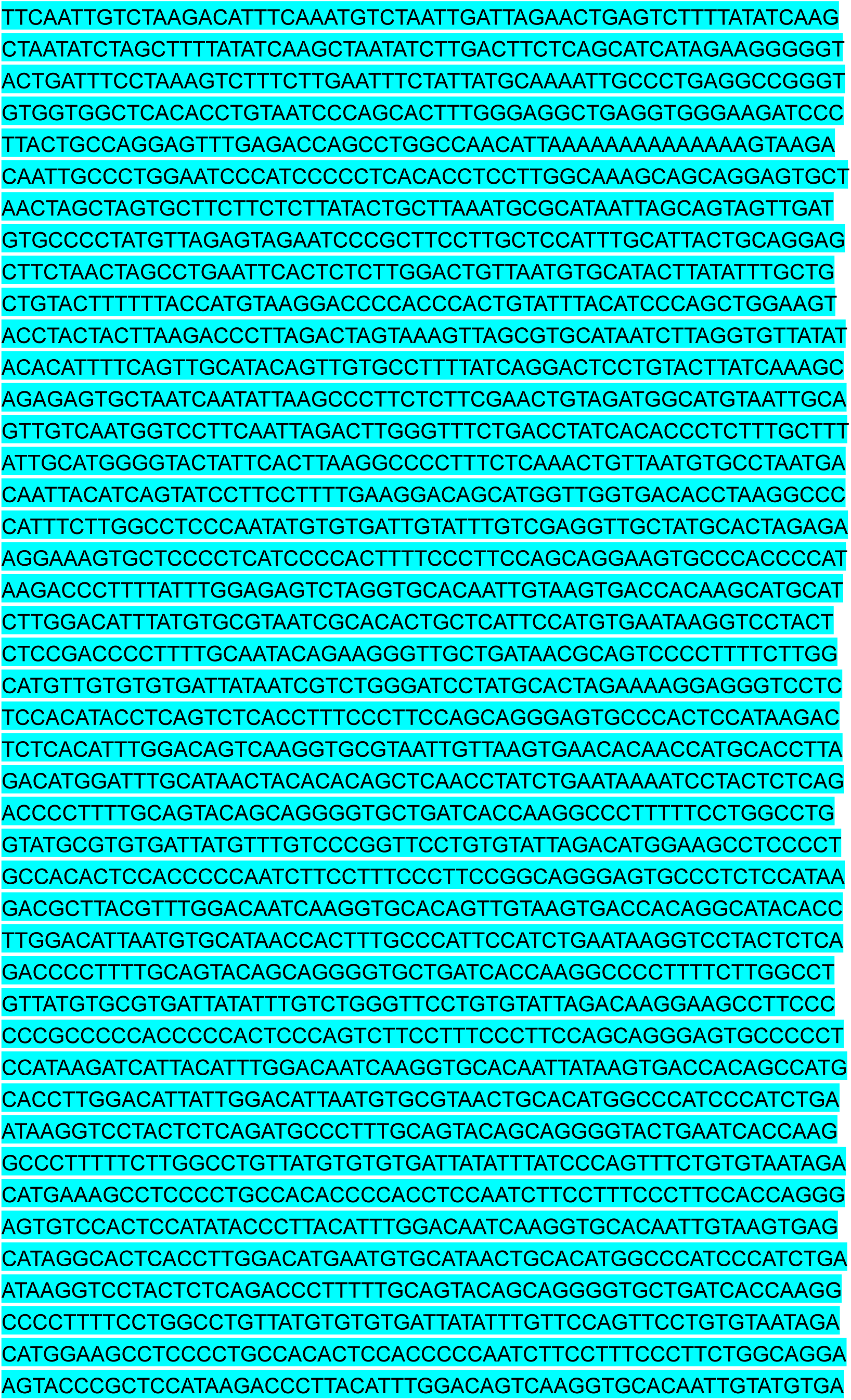
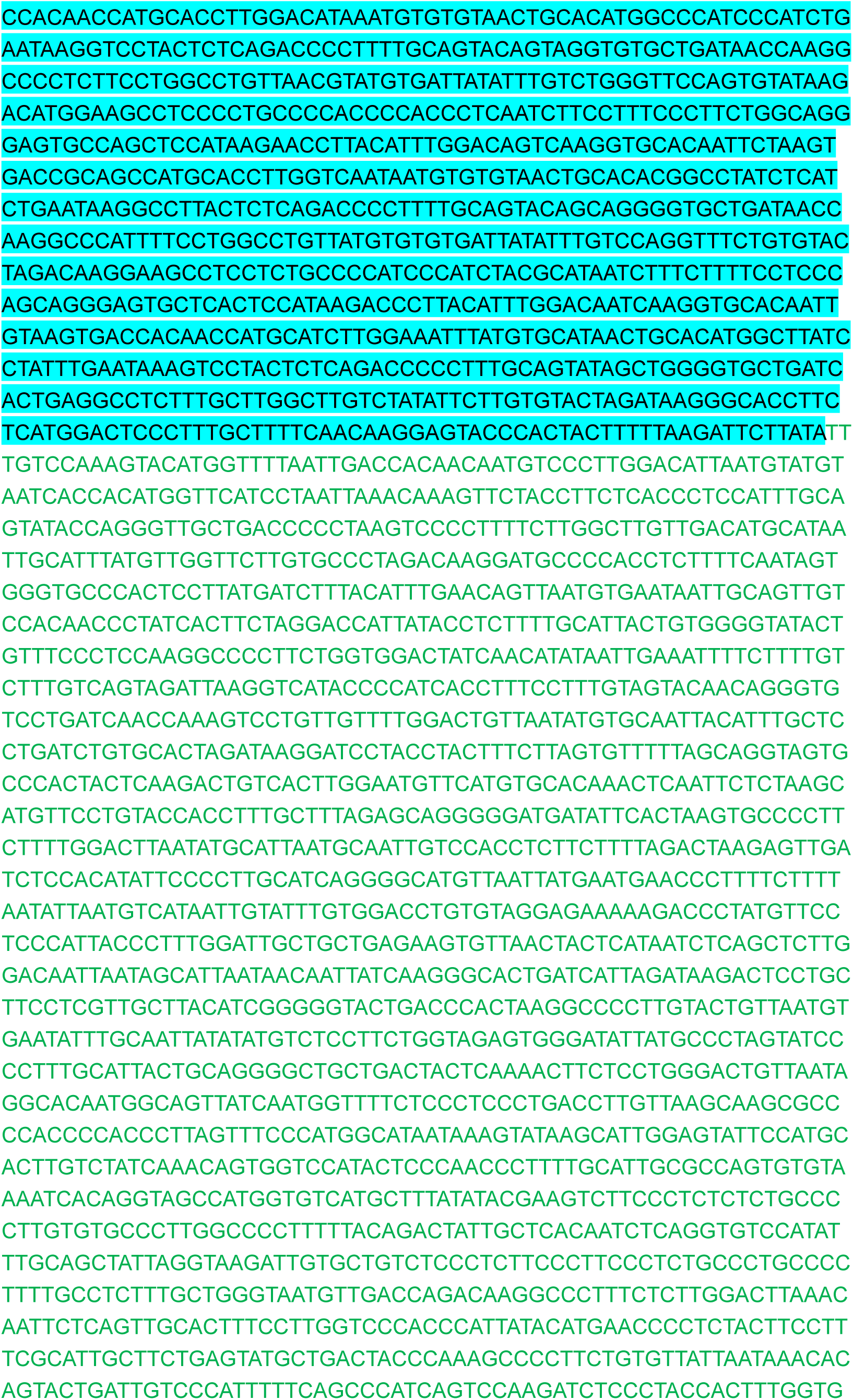
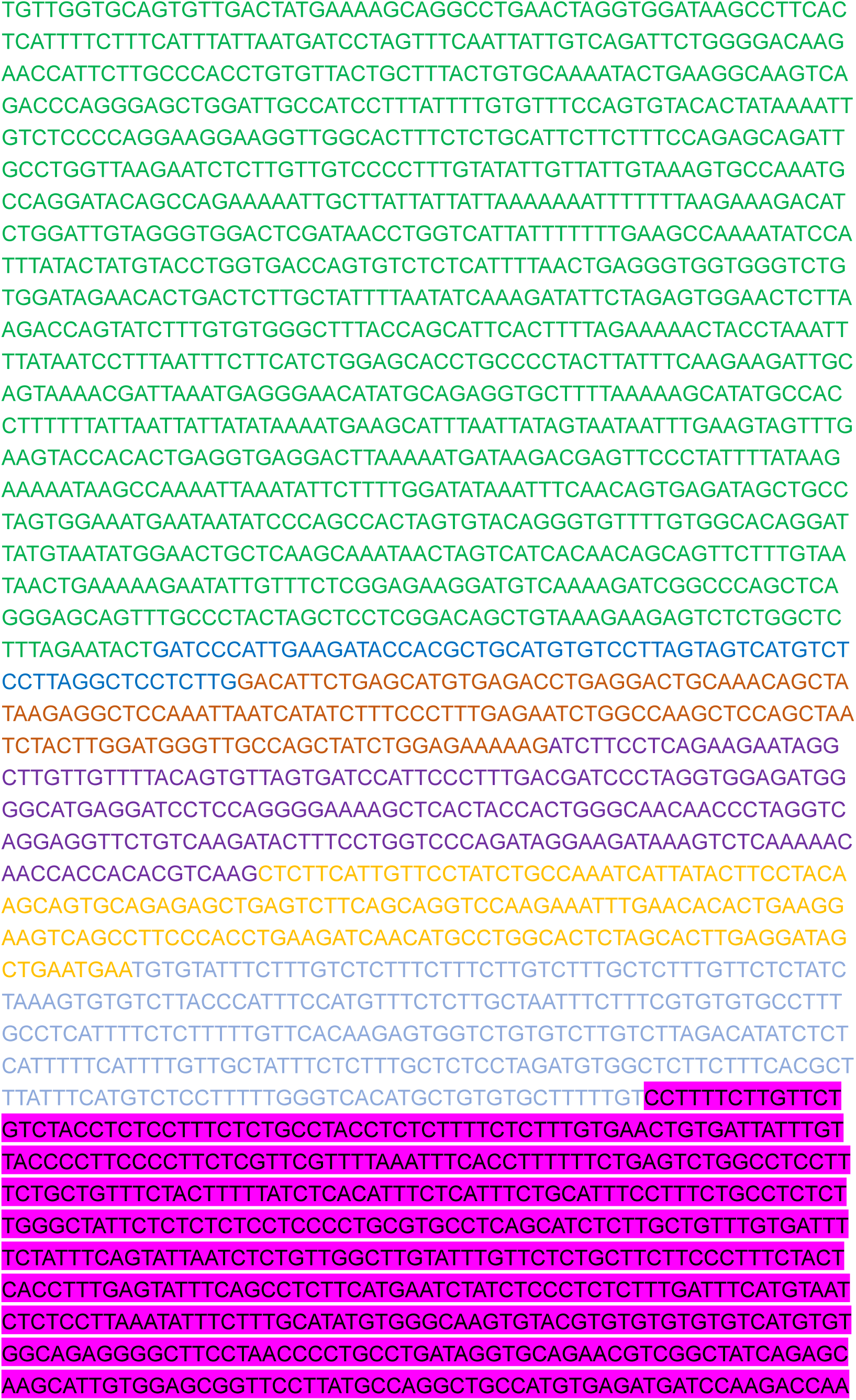
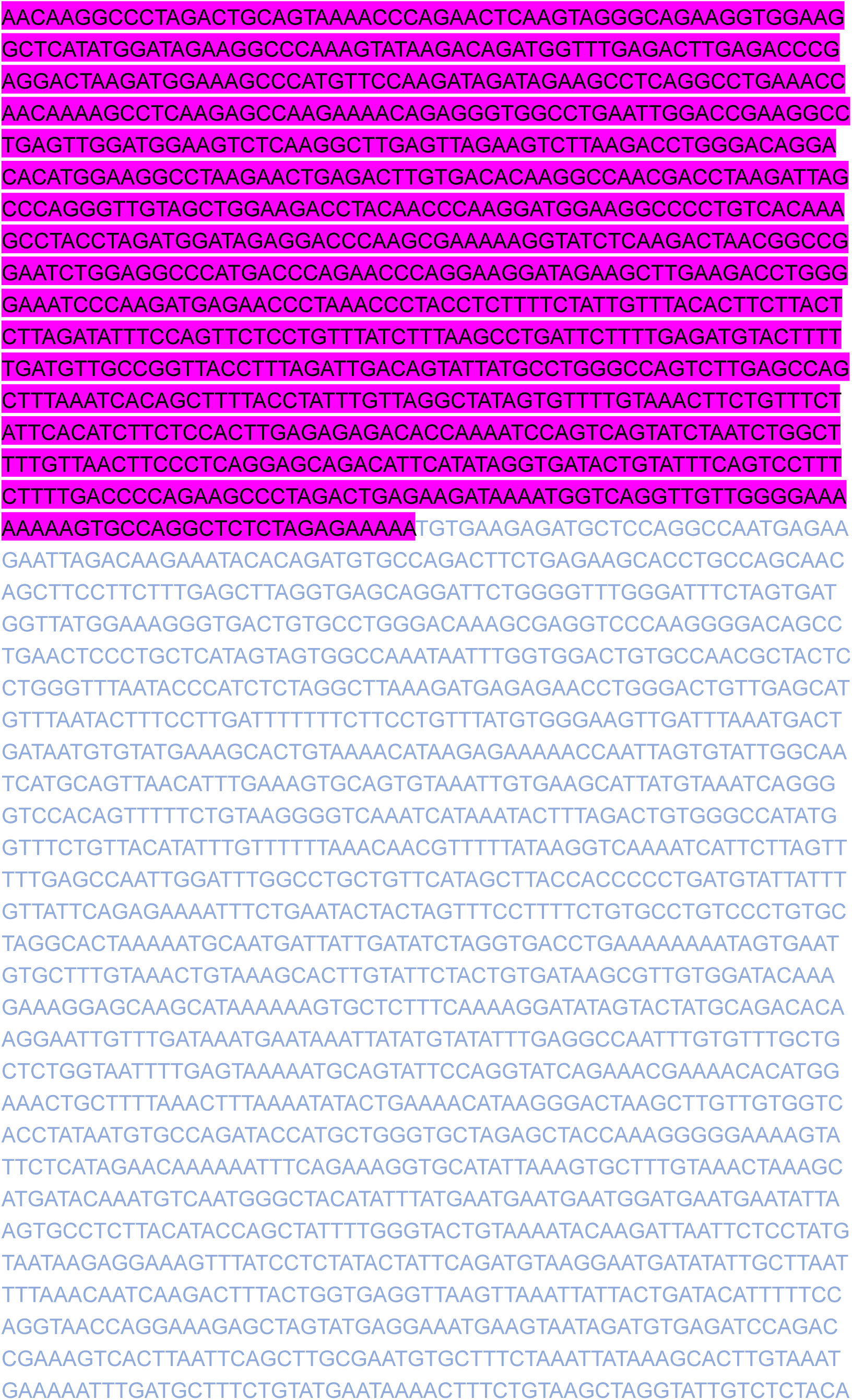
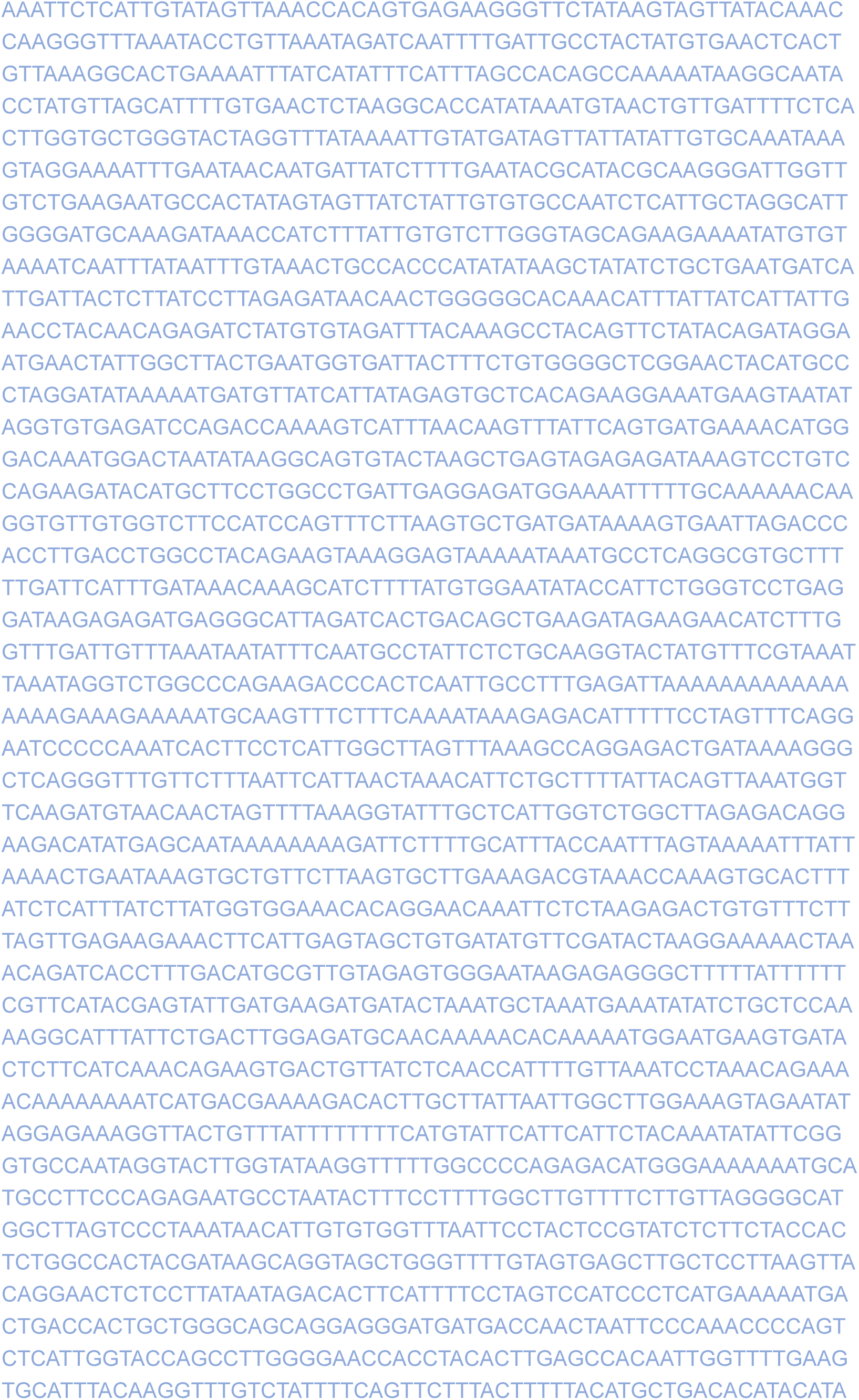
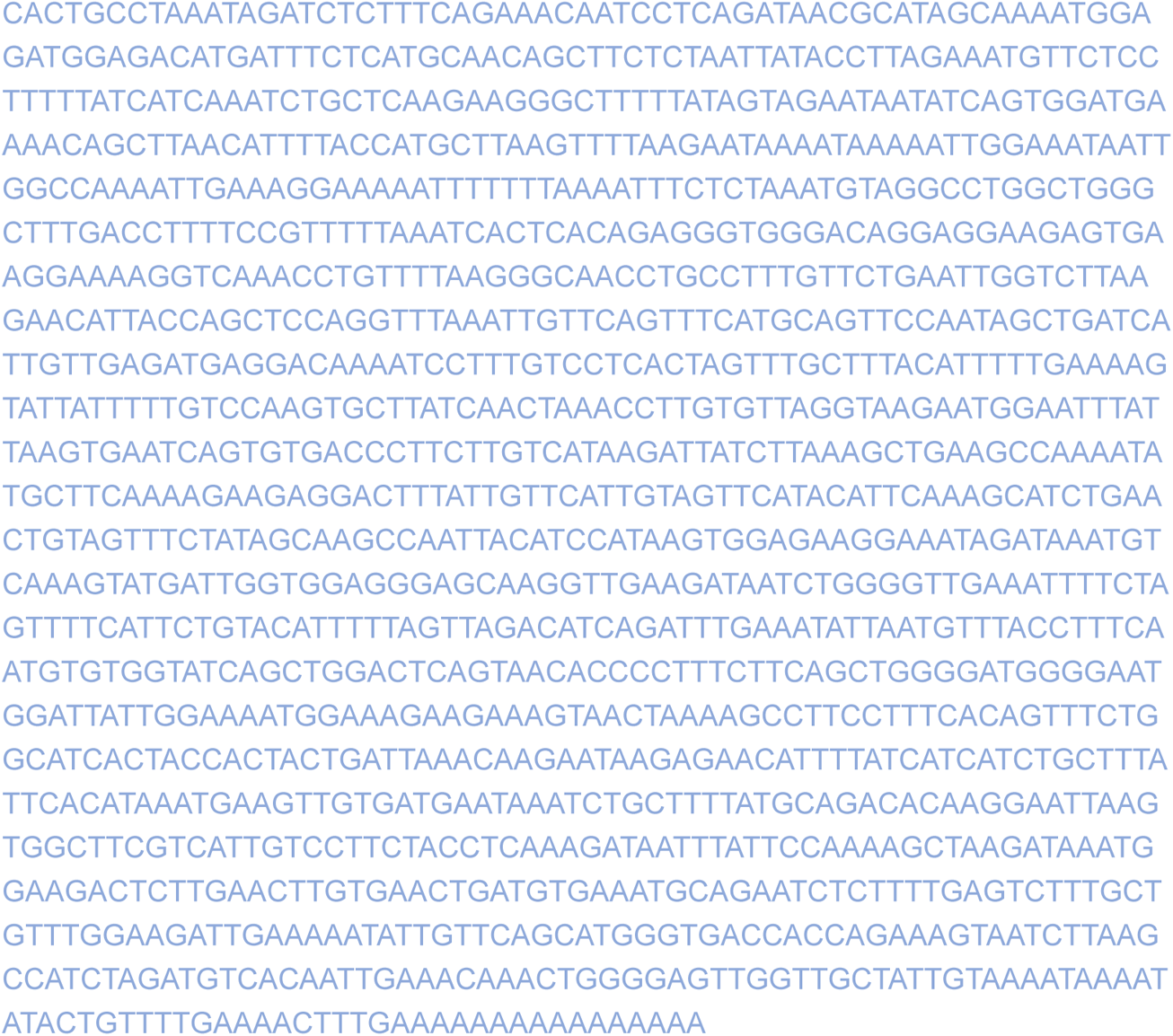
Sequence of the human *XIST* locus.

**Table 6.**
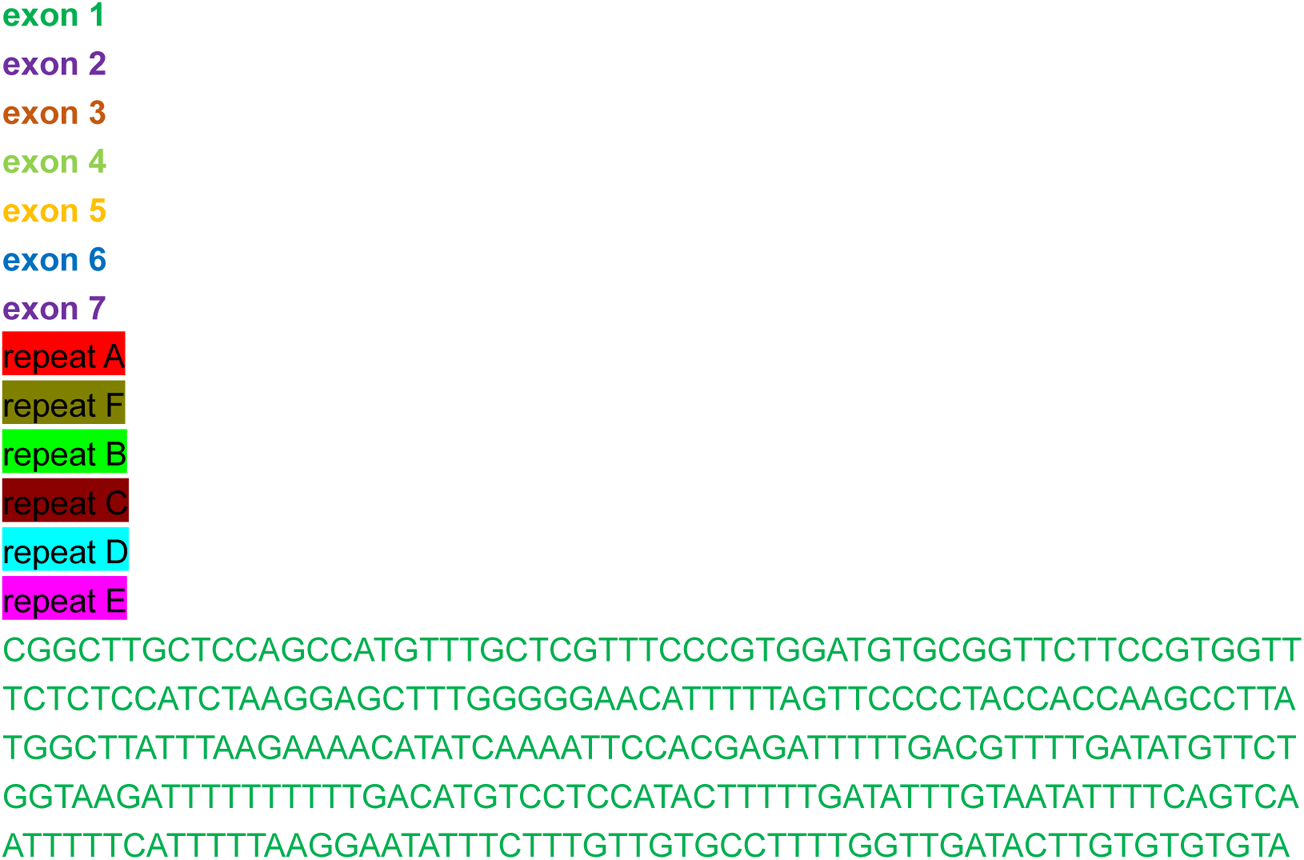
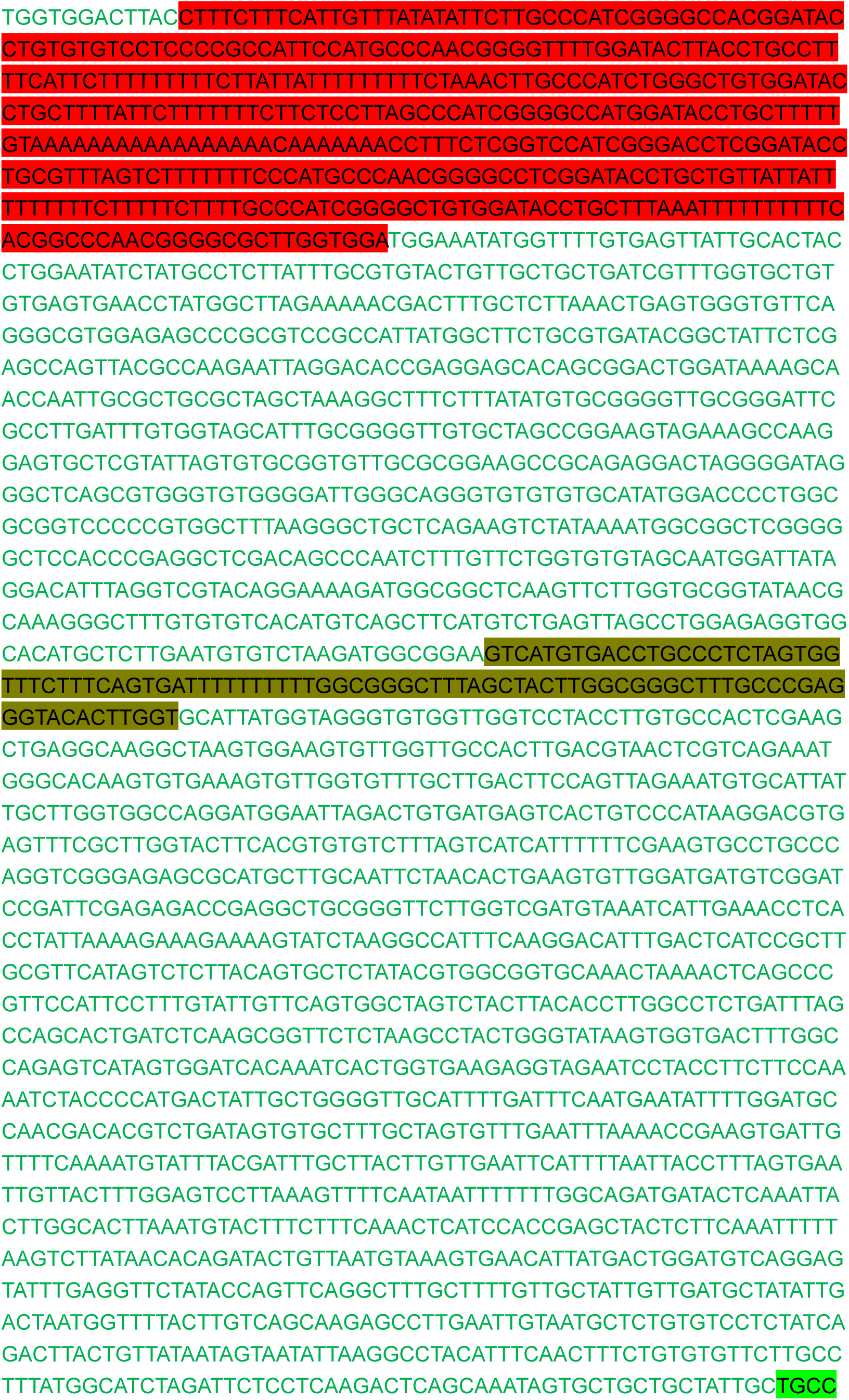
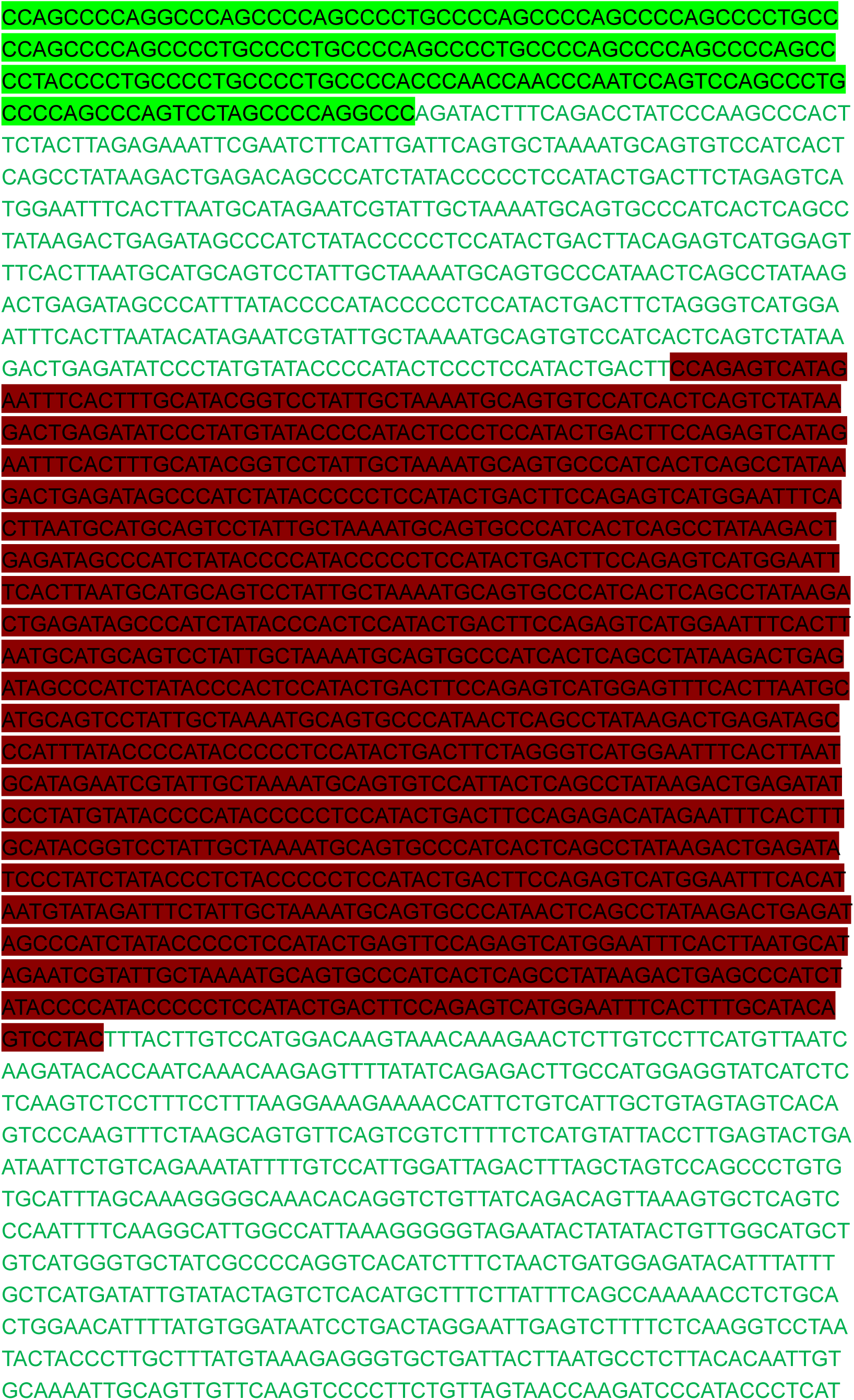
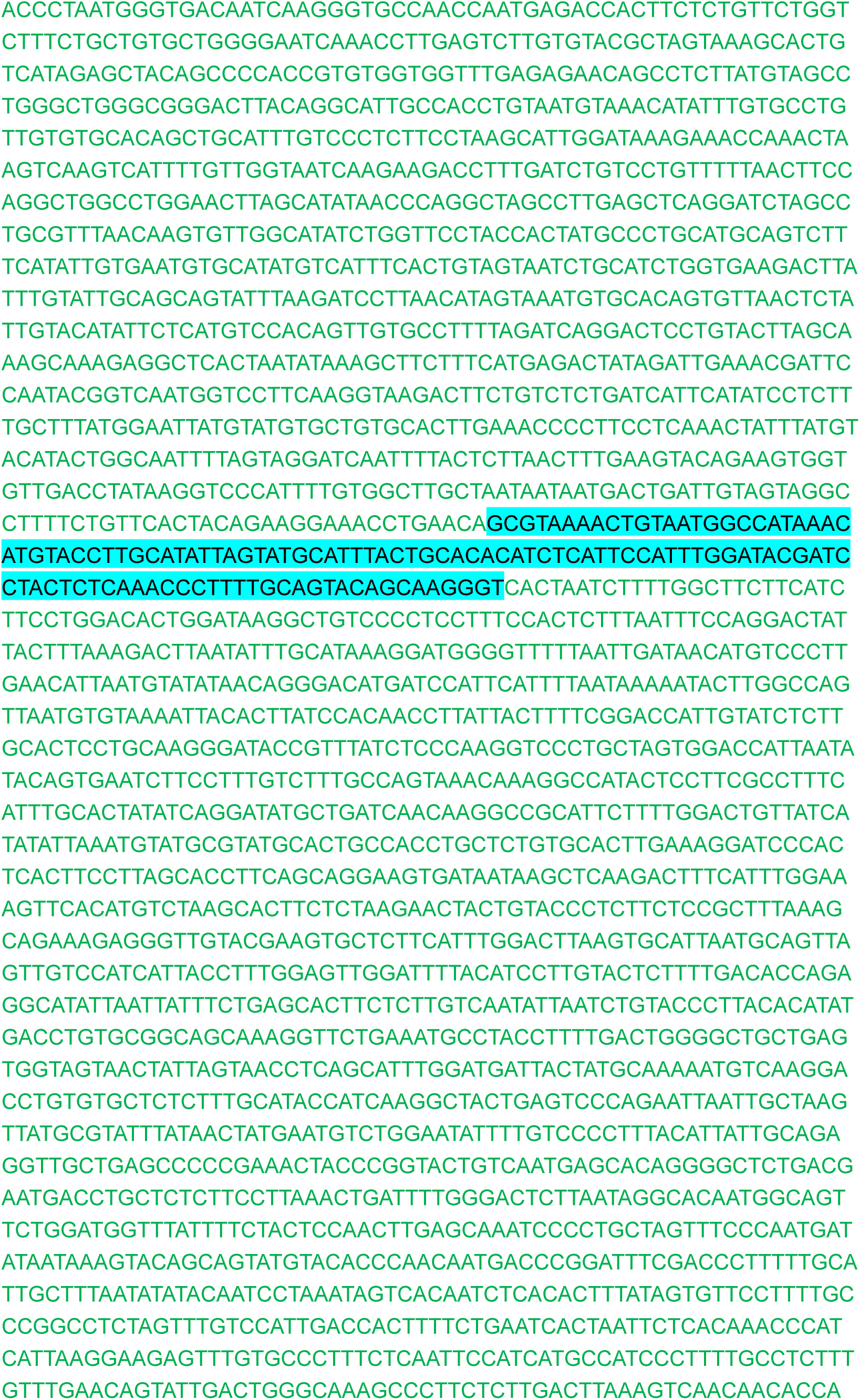
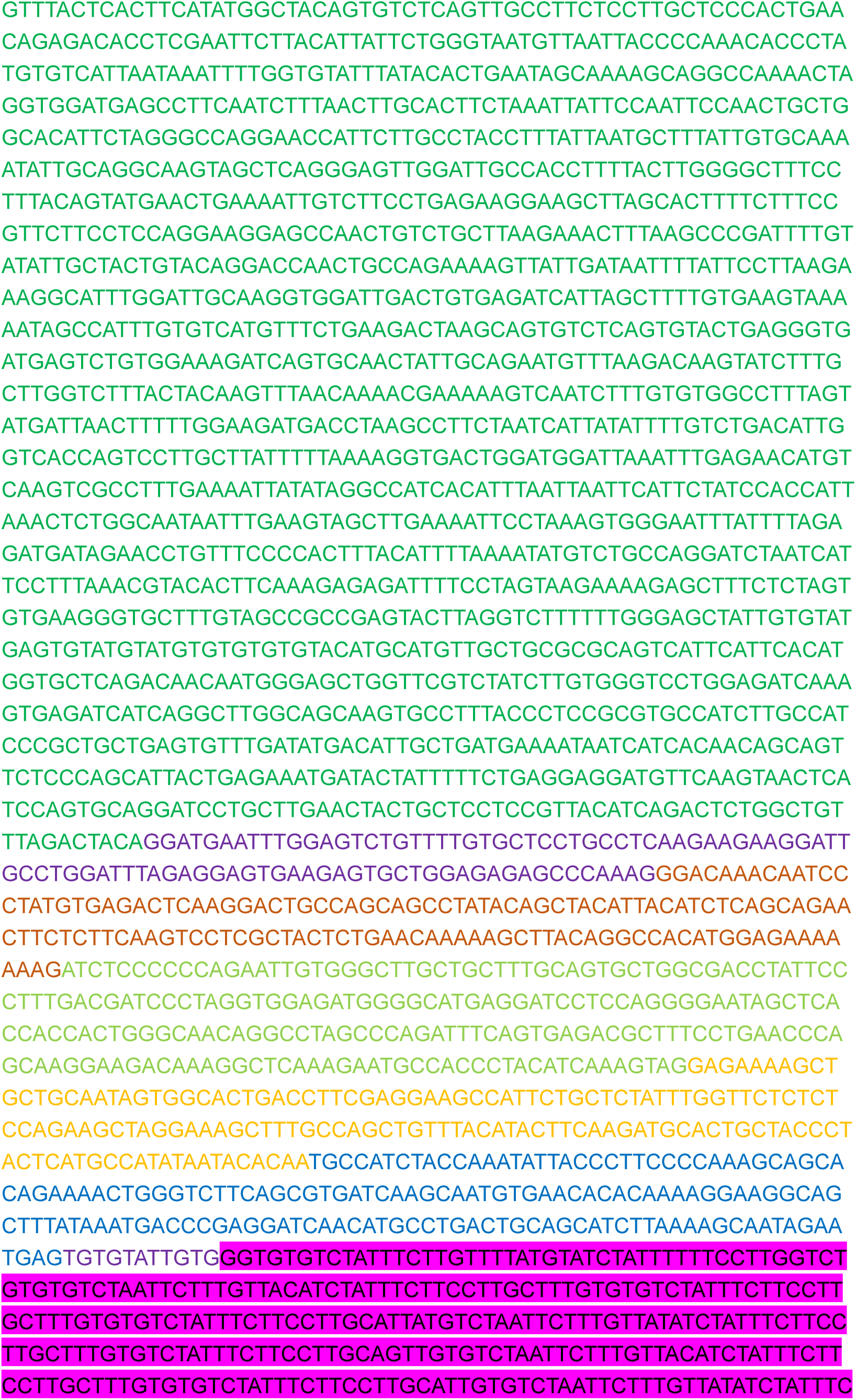
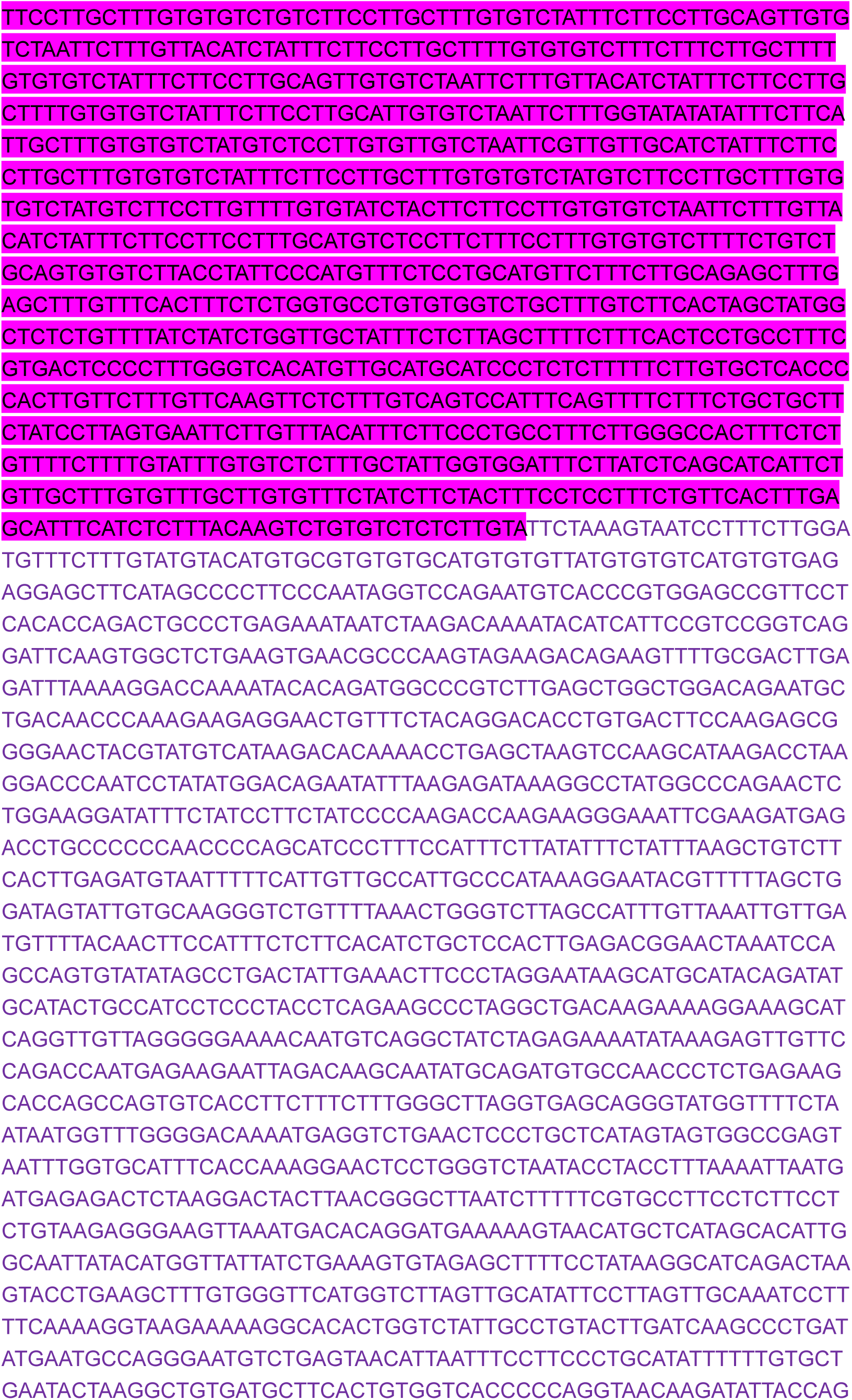
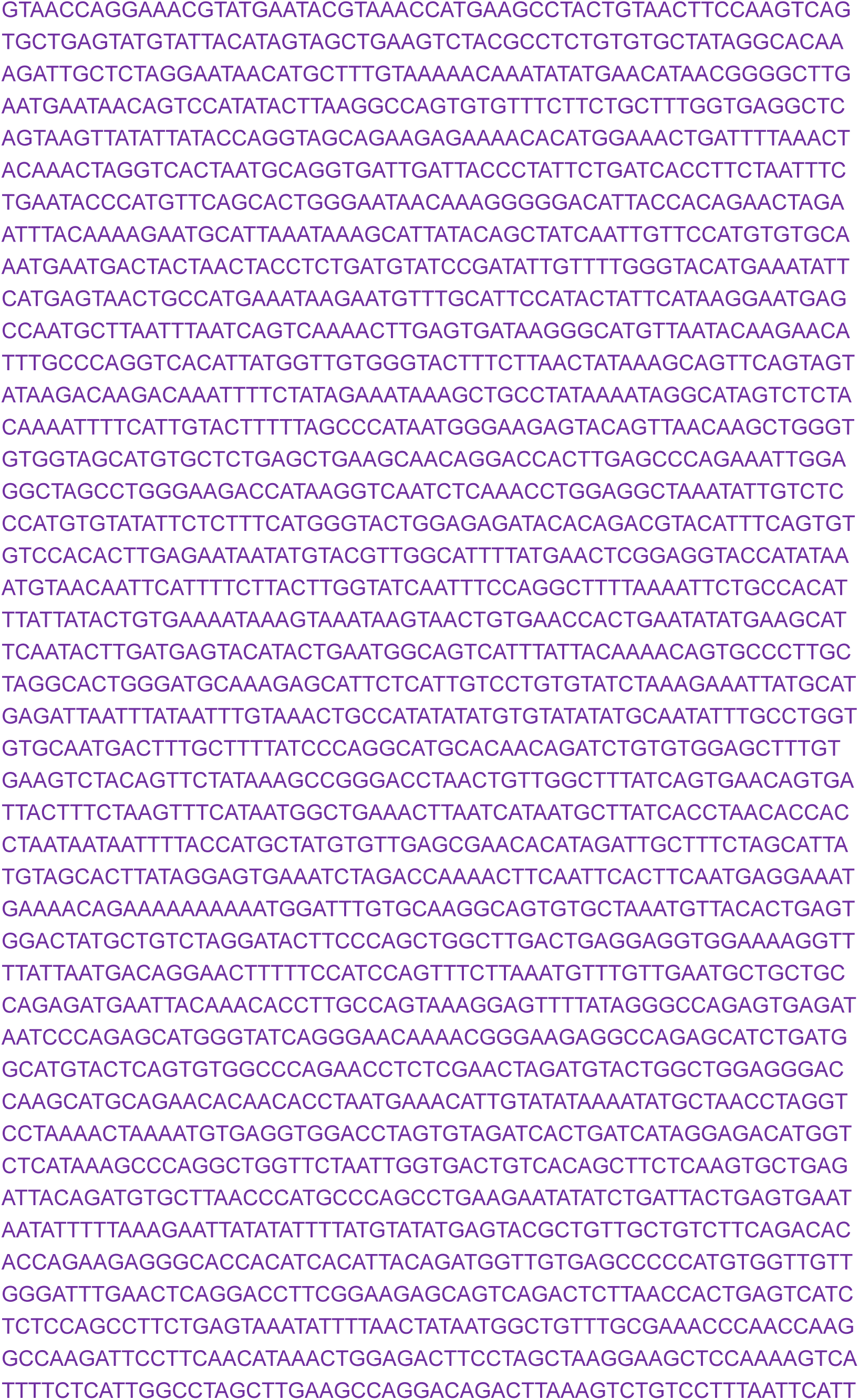
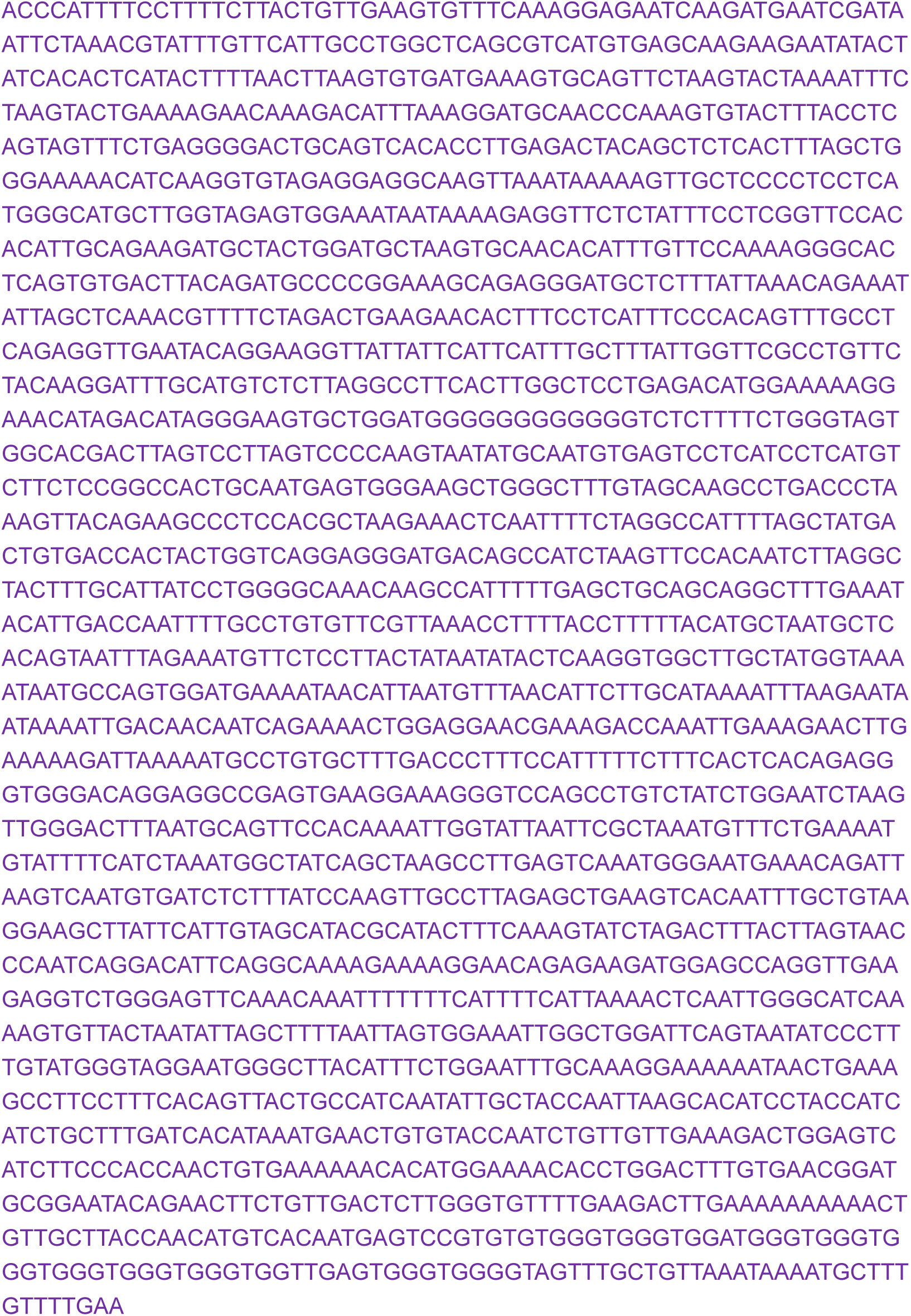
Sequence of the mouse *Xist* locus.

